# Mechanistic insights into the UFM1 E3 ligase complex in ufmylation and ribosome-associated protein quality control

**DOI:** 10.1101/2023.02.16.528878

**Authors:** Ryosuke Ishimura, Sota Ito, Gaoxin Mao, Satoko Komatsu-Hirota, Toshifumi Inada, Nobuo N Noda, Masaaki Komatsu

## Abstract

Ubiquitin-fold modifier 1 (UFM1) is a ubiquitin-like protein covalently conjugated with intracellular proteins through ufmylation, similar to ubiquitylation. Ufmylation is involved in processes such as endoplasmic reticulum (ER)-associated protein degradation, ribosome-associated protein quality control (RQC) at the ER (ER-RQC), and ER-phagy. However, it remains unclear how ufmylation regulates such distinct ER-related functions. Herein, we provide insights into the mechanism of the UFM1 E3 complex in not only ufmylation but also ER-RQC. The E3 complex consisting of UFL1 and UFBP1 interacted with UFC1, UFM1 E2, and subsequently CDK5RAP3, the last of which is an adaptor for ufmylating ribosomal subunit RPL26. When CDK5RAP3 was absent from the E3 complex, UFBP1 ufmylation occurred, a process thought to drive ER-phagy. Further, upon treatment with anisomycin, an inducer of disome formation, the UFM1 E3 complex associated with ufmylated RPL26 on the 60S ribosomal subunit through the UFM1-interacting region of UFBP1. Loss of E3 components or disruption of the interaction between UFBP1 and ufmylated RPL26 attenuated ER-RQC. These results clarify the molecular mechanism of the UFM1 system and provide new insights into the role of ufmylation.

## Introduction

Post-translational modifications amplify limited genomic information and extend the functions of single proteins in diverse cellular processes. Among such modifications, the best characterized are those involving ubiquitin or ubiquitin-like modifiers (UBLs) (*1–4*). Modification systems in which single or multiple ubiquitins or UBLs bind to proteins act as signal converters that regulate various cellular functions. These are catalyzed by a highly regulated and elaborate system within cells, and are carried out by sequential reactions of multiple enzymes comprising an activating enzyme (E1), a conjugating enzyme (E2), and a ligase enzyme (E3). Initially, an E1 forms a high-energy thioester bond with a specific ubiquitin or its corresponding UBLs via adenylation in an ATP-dependent manner. The ubiquitin or UBLs activated by E1 are then transferred to a specific E2 via thioester bonds. In some cases, E2 can directly transfer the ubiquitin or UBLs to substrate proteins in an isopeptide linkage. However, E2s usually require the participation of E3 to achieve substrate-specific ubiquitylation or UBL conjugation in cells. E3 is defined as the enzyme required to recognize the specific substrate and plays a central role in ubiquitination or UBL conjugation (*5*). Indeed, compared with the two E1s and approximately 30 E2s, the number of E3s, which contain a core domain such as RING, HECT, or U-box, is estimated to exceed 600 in humans. Ubiquitination and most UBL conjugation reactions are reversible, with deubiquitinating enzymes removing ubiquitin and UBLs from target proteins and regulating ubiquitination and UBL binding. Covalent modification of intracellular proteins by ubiquitin mediates selective protein degradation, mainly by the 26S proteasome and autophagy, while modification of intracellular proteins by UBLs regulates protein localization, protein-protein and protein-DNA interactions, and biochemical activity.

Ubiquitin-fold modifier 1 (UFM1) is a UBL that is covalently bound to intracellular proteins by ufmylation, a reaction similar to ubiquitination (*6, 7*). UFM1 is synthesized as a pro-form, and the two amino acids at the C-terminus of pro-UFM1 are cleaved by specific proteases, UFSP1 and UFSP2 (*8, 9*). The result is a mature form in which the glycine residues essential for conjugation are exposed. The mature form of UFM1 is activated by forming a high-energy thioester bond with UBA5, a UFM1-specific E1 enzyme. Activated UFM1 is transferred to UFC1, a UFM1-specific E2 enzyme that is covalently bound to the target protein via a UFM1-specific E3 enzyme complex comprising UFL1, UFBP1 (also called DDRGK1 or C20orf116), and CDK5RAP3 (*10–13*). Of these E3 components, UFL1 and UFBP1 are essential for ufmylation (*14, 15*), while CDK5RAP3 may be involved in the regulation of E3 activities such as polyufmylation and direction of ufmylation on the ribosomal protein RPL26 (*13*). This reaction is reversible, as covalently bound UFM1 is cleaved by the cysteine protease UFSP2 (*8, 16*). In mice, disruption of the UFM1 system results in abnormalities in erythroid differentiation (*11, 12, 17, 18*), liver development (*12*), and neurogenesis (*19, 20*). Human genetic mutations in *UFM1, UBA5*, and *UFC1* lead to reduced ufmylation and hereditary paediatric encephalopathy (*19, 21–23*). Mutations in *UFSP2* also cause two autosomal dominant disorders, namely Beukes hip dysplasia (*24*) and vertebral epiphyseal dysplasia (*25*).

UFBP1 possesses a signal peptide for the endoplasmic reticulum (ER) and a transmembrane helix localized on the ER (*10, 26, 27*), and UFL1 forms a stable complex with UFBP1 (*13, 28*), suggesting potential roles of the UFM1 system on the ER. Indeed, the UFM1 system has long been associated with ER stress and ER-associated degradation (*7*). Further, accumulating evidence has recently indicated that ufmylation plays an important role in ribosome-associated protein quality control at the ER (ER-RQC) (*29–31*), as well as in selective autophagy of the ER (ER-phagy) (*27, 32, 33*). The former seems to be mediated by the ufmylation of ribosomal proteins such as RPL26 (*29, 30*), and the latter by the ufmylation of a UFM1 E3 component, UFBP1, and an ER-anchoring protein, CYB5R3 (*10, 27, 34*). However, aside from the X-ray crystal structures of UFM1 (*35*), UBA5 (*36*), UFC1 (*37*), and their complex (*38, 39*), the structure of the UFM1 E3 complex is unknown, and the UFM1 E3 components show no similarity to any previously identified E3 of ubiquitin or UBLs, making it difficult to infer their structure. Therefore, the molecular mechanism(s) by which the UFM1 E3 complex regulates diverse functions on the ER, such as ER-RQC and ER-phagy, is largely unknown.

Herein, we provide insights into the mechanism of the UFM1 E3 ligase complex in ufmylation and its association with the 60S ribosomal subunit through biochemical analyses and structure prediction using AlphaFold2 (AF2) (*40, 41*) with the AlphaFold-Multimer mode (*42*). Further, we reveal that upon treatment with anisomycin, an inducer of disome formation, the UFM1 E3 complex binds to ufmylated RPL26 on the 60S ribosomal subunit through the UFM1-interacting region of UFBP1, which is indispensable for ER-RQC. Our findings indicate that the UFM1 E3 complex is a ligase and also acts as an essential factor for ER-RQC.

## Results

### Prediction of the UFL1-UFBP1-CDK5RAP3 E3 complex structure

Highly accurate protein structure prediction by AlphaFold2 (AF2) is radically changing life science research. Although the structures of the three components comprising the E3 ligase for ufmylation (UFL1, UFBP1, and CDK5RAP3) have never been determined experimentally, their predicted structures are available in the AlphaFold Protein Structure Database (Fig. S1) (*43*). UFL1 comprises two N-terminal α-helices, five tandem winged-helix (WH) domains, and a C-terminal helical domain that contains an intrinsically disordered region (IDR). In contrast, UFBP1 is composed of one N-terminal transmembrane helix, an IDR containing a coiled coil, one WH domain, and two C-terminal α-helices. CDK5RAP3 consists mainly of α-helices: two α-helices (residues 109–168 and 427–493) together form an anti-parallel coiled coil that is as long as ~10 nm, and two globular domains are located at both ends of the coiled coil, leading to a dumbbell-like shape. One of the globular domains is composed of ~120 N-terminal residues and ~30 C-terminal residues, and is thus named the terminal dumbbell (T-dumbbell), whereas the other globular domain is named the central dumbbell (C-dumbbell). The C-dumbbell possesses an IDR containing 80 residues.

Although AF2 was initially not good at predicting the structures of heterocomplexes, the recent implementation of the AlphaFold-Multimer mode in AF2 (*42*) now enables us to predict these structures with much greater accuracy. Using AF2 with the AlphaFold-Multimer mode (hereafter, all the AF2 predictions that are mentioned were performed using this mode), we predicted the structure of the UFL1-UFBP1-CDK5RAP3 E3 complex. To further increase the prediction accuracy, we initially used truncated forms of UFL1 (residues 1–302) and UFBP1 (residues 209–314) and obtained a well-converged predicted model of the UFL1(1–302)-UFBP1(209–314)-CDK5RAP3 complex (Fig. S2A). After that, we superimposed the full-length AF2 structures of UFL1 and UFBP1 on each truncated form and obtained the full-length UFL1-UFBP1-CDK5RAP3 E3 complex model (Fig. 1A). Upon complex formation, the second N-terminal α-helix of UFL1 and the two C-terminal α-helices of UFBP1 fold into one WH domain, forming a complex with seven consecutive WH domains. The UFBP1 WH domain is named WH1, the five WH domains of UFL1 are named WH3 to WH7 (counted from the N-terminus), and the WH domain created upon complex formation is named WH2. These predictions are consistent with the structure of the UFL1-UFBP1 complex recently predicted using AF2 (*13, 28*). In the predicted ternary complex, CDK5RAP3 interacts with the six tandem WH domains (WH1-WH6) of the UFL1-UFBP1 complex and the N-terminal helix of UFL1 so that the long axis of CDK5RAP3 and the WH1-WH6 portion of the UFL1-UFBP1 complex become approximately parallel (Fig. 1 A).

**Figure 1.**
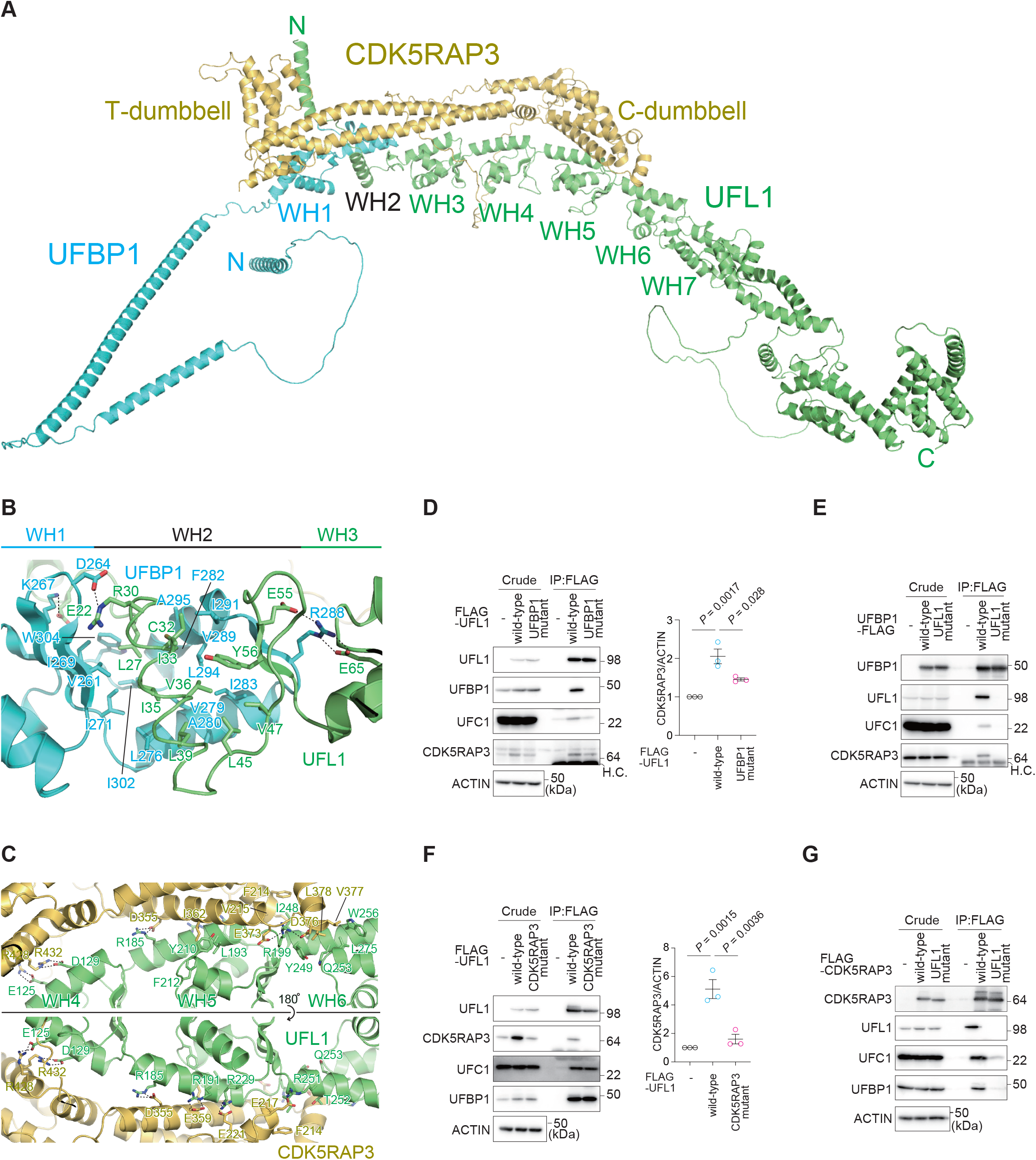
Structure and interaction modes of the UFM1 E3 complex. **(A)** Predicted three-dimensional structure of the full-length UFL1-UFBP1-CDK5RAP3 E3 complex. **(B, C)** Binding mode between UFL1 and UFBP1 (B) and between UFL1 and CDK5RAP3 (C). The side chains involved in interactions are shown with a stick model, where oxygen and nitrogen atoms are colored red and blue, respectively. Broken lines indicate possible salt bridges. **(D-G)** Immunoprecipitation assay. FLAG-tagged wild-type UFL1 and UFBP1-interaction defective UFL1 mutant (UFL1^UFBP1 mutant^) were transfected into *UFL1*-deficient HEK293T cells (D). FLAG-tagged wild-type UFBP1 and UFL1-interaction defective UFBP1 (UFBP1^UFL1 mutant^) were transfected into *UFBP1*-deficient HEK293T cells (E). FLAG-tagged wild-type UFL1 and CDK5RAP3-interaction defective UFL1 mutant (UFL1^CDK5RAP3 mutant^) were transfected into *UFL1*-deficient HEK293T cells (F). FLAG-tagged wild-type CDK5RAP3 and UFL1-interaction defective CDK5RAP3 (CDK5RAP3^UFL1 mutant^) were transfected into *CDK5RAP3*-deficient HEK293T cells (G). Forty-eight hours after transfection, cells were lysed and immunoprecipitated with anti-FLAG-M2 gel, and the immunoprecipitants were then subjected to immunoblot analysis with the indicated antibodies. Bar graph shows the results of quantitative densitometric analysis of CDK5RAP3 relative to actin (*n* = 3) (F). Data are means ± s.e. Statistical analysis was performed by Welch’s *t*-test. Data shown are representative of three separate experiments.

### Interactions leading to construction of the UFM1 E3 complex

As described above, UFL1-UFBP1 interaction results in creation of the WH2 domain, which is composed of three α-helices (one from UFL1 and two from UFBP1) and a three-stranded β-sheet (two strands from UFL1 and one from UFBP1). In detail, Cys32, Ile33, Val36, Leu39, Leu45, Val47, and Tyr56 of UFL1 form hydrophobic interactions with Leu276, Val279, Ala280, Ile283, Ile291, Leu294, and Ala295 of UFBP1, and these interactions form the hydrophobic core of the WH2 domain (Fig. 1B). The WH2 domain closely interacts with the WH1 domain via hydrophobic interactions formed between Leu27 and Ile35 of UFL1 and Val261, Ile269, Ile271, Ile302, and Trp304 of UFBP1. In addition to hydrophobic interactions, four possible salt-bridge pairs are observed between UFL1 (Glu22, Arg30, Glu55, Glu65) and UFBP1 (Asp264, Lys267, Arg288). Besides these, dozens of hydrogen bonds are formed between UFL1 and UFBP1. These hydrophobic and hydrophilic interactions bury a surface area of 1350 Å^2^ for each protein.

Both the C- and T-dumbbells of CDK5RAP3 interact with the UFL1-UFBP1 complex. The T-dumbbell binds to the WH4, WH5, and WH6 domains of UFL1 (Fig. 1C). Eight possible salt-bridge pairs are observed between UFL1 (Glu125 and Asp129 from WH4; Arg185, Arg191, Arg199, and Arg229 from WH5; Arg251 from WH6) and CDK5RAP3 (Glu217, Glu221, Asp355, Glu359, Glu373, Asp376, Arg428, Arg432). Besides these, dozens of hydrogen bonds are formed between UFL1 and the T-dumbbell. In addition, several hydrophobic interactions are observed, including those between Ile248, Tyr249, Thr252, Gln253, Trp256, and Leu275 of the WH6 domain and Phe214, Val215, Val377, and Leu378 of CDK5RAP3, and between Leu193, Tyr210, and Phe212 of WH5 and Ile362 of CDK5RAP3. These interactions bury a surface area of 1780 Å^2^ for each protein. In addition to the C-dumbbell-mediated interactions, CDK5RAP3 uses the T-dumbbell to bind to the N-terminal helix of UFL1 and the WH1 domain of UFBP1, mainly through hydrophilic interactions, which buries surface areas of 630 and 490 Å^2^, respectively (Fig. S2B). These observations suggest that CDK5RAP3 binds to the UFL1-UFBP1 complex mainly via recognition of UFL1 WH4-6 by the C-dumbbell.

### UFL1 plays a central role in E3 complex formation

To confirm the binding mode biochemically, we constructed UFL1 and UFBP1 mutants in which amino acids involved in the hydrophobic interactions were replaced with Ala (UFL1^L27A C32A I35A L39A^ and UFL1^UFBP1^ mutant; UFBP1^I271A L276A I302A W304A^ and UFBP1^UFL1 mutant^). To exclude the effect of endogenous UFL1, we generated *UFL1* -deficient HEK293T cells (Fig. S3A). FLAG-tagged wild-type UFL1 and UFL1^UFBP1 mutant^ were expressed in *UFL1*-deficient HEK293T cells, and we verified that the expression levels were similar (Fig. 1D). An immunoprecipitation assay using anti-FLAG antibody revealed that wild-type UFL1 but not UFL1^UFBP1 mutant^ interacted with endogenous UFBP1 (Fig. 1D). In addition to UFBP1 and CDK5RAP3 (*10–13*), UFL1 directly interacts with UFM1 E2, UFC1 (*10*). The immunoprecipitant from cells expressing the UFL1^UFBP1 mutant^ contained endogenous UFC1 and CDK5RAP3, but the levels of both were less than those from wild-type UFL1-expressing cells (Fig. 1D). FLAG-tagged wild-type UFBP1 and UFBP1^UFL1 mutant^ were expressed in *UFBP1* knockout (KO) HEK293T cells (*27*), and the cell lysates were immunoprecipitated with anti-FLAG antibody. The immunoprecipitant prepared from wild-type UFBP1-expressing cells contained endogenous UFL1, UFC1, and CDK5RAP3 (Fig. 1E). UFL1, UFC1, and CDK5RAP3 were not detected in the immunoprecipitants of cells expressing UFBP1^UFL1 mutant^ (Fig. 1E).

In the next series of experiments, to biochemically investigate the interaction of UFL1 with CDK5RAP3, we constructed UFL1 and CDK5RAP3 mutants in which the amino acids involved in the electrostatic interaction between the two proteins were substituted with Ala (UFL1^D129A R185A R191A R199A R251A^ and UFL1^CDK5RAP mutant^; CDK5RAP3^E217A D355A E359A E373A R432A^ and CDK5RAP3^UFL1 mutant^). FLAG-tagged wild-type UFL1 and UFL1^CDK5RAP mutant^ were expressed in *UFL1*^-/-^ HEK293T cells, and we verified that both wild-type and mutant UFL1 were expressed at similar levels (Fig. 1F). Immunoprecipitation with anti-FLAG antibody showed that wild-type UFL1, but not UFL1^CDK5RAP mutant^, bound to endogenous CDK5RAP3 (Fig. 1F). Like wild-type UFL1, the UFL1 mutant showed the ability to bind to UFC1 and UFBP1 (Fig. 1F). To investigate the effect of the CDK5RAP3 mutant on the interaction with UFL1, we utilized *CDK5RAP3* KO HEK293T cells (*27*) and expressed FLAG-tagged wild-type CDK5RAP3 and CDK5RAP3^UFL1 mutant^. The cell lysates were immunoprecipitated with anti-FLAG antibody, and the immunoprecipitants were subjected to immunoblot analysis with anti-UFL1 and anti-CDK5RAP3 antibodies. The expression levels of wild-type CDK5RAP3 and the mutant were comparable (Fig. 1G). The immunoprecipitation assay showed that wild-type but not mutant CDK5RAP3 interacted with endogenous UFL1 (Fig. 1G). The immunoprecipitants from cells expressing the CDK5RAP3 mutant contained hardly any UFC1 or UFBP1 (Fig. 1G). These results indicate that UFL1 plays a central role in the interactions between UFC1, UFBP1, and CDK5RAP3.

### Structural basis of UFC1-UFL1 interaction

E3 promotes ubiquitylation-like conjugation reactions through direct interaction with E2. We next used AF2 to predict how E3 components recognize UFC1. Structural prediction of the full-length UFL1-UFC1 binary complex suggests that UFC1 is mainly recognized by the N-terminal α-helix (residues 1–21) of UFL1, which we hereafter refer to as the UFC1-binding sequence (UBS) (Fig. S4A). All of the top five prediction models show similar interaction between UFL1^UBS^ and UFC1, although the relative orientation between UBS and the other region of UFL1 is variable (Fig. S4B). Structural prediction of the UFL1^UBS^-UFC1 binary complex gives us almost the same interaction model (Fig. 2A, S4C). These predictions are consistent with our previous report that UFL1 directly recognizes UFC1 (*10*). In addition, AF2 predicted direct interaction of UFBP1 with UFC1 (the best model is shown in Fig. S5D); however, the top five prediction models each have distinct UFBP1-UFC1 interactions (Fig. S4E), suggesting that these models are unreliable. Structural prediction of the UFL1-UFBP1-UFC1 ternary complex provides us with a combined model of the UFL1-UFC1 and UFBP1-UFC1 complexes (Fig. 2B, S4F) in which UFL1-UFC1 and UFBP1-UFC1 interactions bury surface areas of 961 and 555 Å^2^ for each protein, respectively. These structural predictions suggest that UFL1^UBS^ plays a central role in recruiting E2 to the E3 complex.

**Figure 2.**
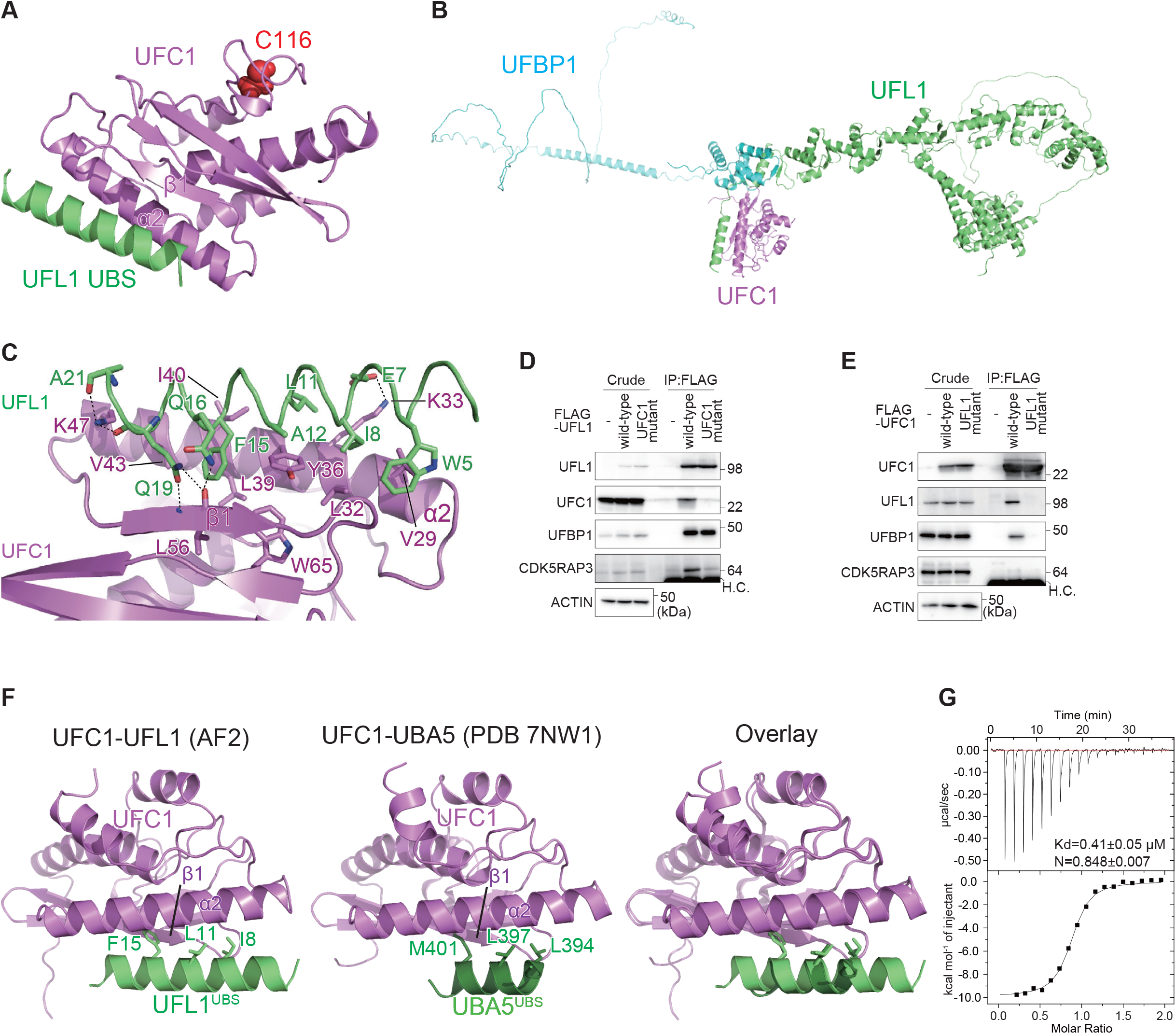
Structure of the UFM1 E3 component UFL1 and the E2 UFC1 complex. **(A)** Structural prediction of the UFL1^UBS^-UFC1 binary complex. Catalytic Cys116 of UFC1 is shown with a space-filling model. **(B)** Structural prediction of the UFL1-UFBP1-UFC1 ternary complex. **(C)** Binding mode between UFL1^UBS^ and UFC1. The side chains involved in interaction are shown with a stick model, where oxygen and nitrogen atoms are colored red and blue, respectively. Broken lines indicate possible electrostatic interactions. **(D, E)** Immunoprecipitation assay. FLAG-tagged wild-type UFL1 and UFC1-interaction-defective UFL1 mutant (UFL1^UFC1 mutant^) were transfected into *UFL1*-deficient HEK293T cells (D). FLAG-tagged wild-type UFC1 and UFL1-interaction-defective UFC1 (UFC1^UFL1 mutant^) were transfected into *UFC1*-deficient HEK293T cells (E). Forty-eight hours after transfection, cells were lysed and immunoprecipitated with anti-FLAG-M2 gel; the immunoprecipitants were then subjected to immunoblot analysis with the indicated antibodies. Data shown are representative of three separate experiments. **(F)** Structural comparison of the UFL1^UBS^-UFC1 (left) and UBA5^UBS^-UFC1 (middle) complexes. The side chains of the three hydrophobic residues of UFL1^UBS^ and UBA5^UBS^ important for UFM1 binding are shown with a stick model. The right UBS panel shows the superimposition of the two complexes. **(G)** Binding affinity of UFL1^UBS^ to UFC1 was measured by ITC.

Detailed interaction between UFL1^UBS^ and UFC1 is shown in Fig. 2C. Previous crystallographic studies revealed that UFC1 is composed of three β-strands (β1-β3), four α-helices (α1-α4), and one 3_10_ helix (*37*), which together fold into a canonical E2 fold, although this fold has an additional α-helix at the N-terminus and lacks the two C-terminal α-helices observed in other E2 proteins. The predicted structure of UFC1 alone is essentially similar to the crystal structure except for the six C-terminal residues and the predicted structure of UFC1 bound to UFL1 (Fig. S4G). This suggests that the prediction of the UFC1 structure is accurate in principle and that UFL1 binding has little effect on UFC1 folding. UFL1^UBS^ is bound to α2 and β1 of UFC1, which are located at opposite sides of the catalytic cysteine (Cys116); as such, UFL1^UBS^ has no apparent effect on the catalytic site structure or its accessibility (Fig. 2A). The UFL1^UBS^-UFC1 interaction arises mainly from hydrophobic interactions (Fig. 2C). The side chains of Trp5, Ile8, Leu11, Ala12, and Phe15 of UFL1 interact with those of Val29, Leu32, Tyr36, Leu39, Ile40, and Val43 of UFC1. In addition, UFL1 Glu7 forms a salt bridge with UFC1 Lys33, and the side chains of UFL1 Gln16 and Gln19 form three hydrogen bonds with the main chain of UFC1 Leu56. These interactions bury a surface area of 770 Å^2^ for each protein.

### Interaction of UFC1 with the UFL1-UFBP1 complex is required for CDK5RAP3-recruitment to the E3 complex

To verify the interaction mode of UFL1 with UFC1 biochemically, we constructed UFL1 and UFC1 mutants, which amino acids involved in the hydrophobic interactions of each other were substituted with Ala (UFL1^W5A E7A E16A E19A F15A^, UFL1^UFC1 mutant^ and UFC1^K33A Y36A I40A^, UFC1^UFL1 mutant^). FLAG-tagged wild-type UFL1 or UFL1^UFC1 mutant^ was expressed in the *UFL1* KO cells, and each cell lysates were subjected to immunoprecipitation assay with anti-FLAG antibody followed by immunoblots with anti-UFL1, anti-UFC1, anti-UFBP and anti-CDK5RAP3 antibodies. Both wild-type and mutant UFL1 were expressed at similar level in the *UFL1*-deficient cells (Fig. 2D). As expected, wild-type UFL1 but not mutant UFL1 interacted with endogenous UFC1 (Fig. 2D). While UFL1^UFC1 mutant^ has an ability to bind to UFBP1, it showed less binding affinity with CDK5RAP3 (Fig. 2D). Next, we expressed FLAG-tagged wild-type UFC1 or UFC1^UFL1 mutant^ in *UFC1* KO HEK293T cells (*23*). As shown in Fig. 2E, the expression levels of wild-type and mutant UFC1 was comparable. The immunoprecipitant prepared from the *UFC1*-deficient cells expressing wild-type UFC1 contained endogenous UFL1 (Fig. 2E), but not in the case of the mutant expressing cells (Fig. 2E). Endogenous CDK5RAP3 was present in immunoprecipitants of cells expressing wild-type UFC1 but not the UFL1-interaction defective UFC1 mutants (Fig. 2E). Taken together, these results suggest that the interaction of UFL1 with UFC1 is required for the association of UFL1 with CDK5RAP3.

### Structural mechanism of switching from the E1-E2 complex to the E2-E3 complex

Analysis of the crystal structure of UFC1 complexed with the UBA5^UBS^ suggests that UFL1 and UBA5 interact with UFC1 in very similar ways (Fig. 2F) (*39*). In both UBSs, one α-helix binds to α2 and β1 of UFC1, mainly through hydrophobic interactions. Binding to UFC1 is quite similar between the side chains of L394, L397, and M401 of UBA5^UBS^ and those of I8, L11, and F15 of UFL1^UBS^. The helix of UBA5^UBS^ is shorter than that of UFL1^UBS^ and thus has fewer interactions with UFC1, which is partly compensated for by the interactions mediated by the N-terminal loop of the UBS helix (*39*). Since there is complete overlap between UFL1 and UBA5 in terms of their binding sites on UFC1 (Fig. 2F), it is obvious that E1 (UBA5) and E3 (UFL1-UFBP1) enzymes compete with each other for E2 (UFC1) binding. This competitive binding is also conserved in other Ub/Ubl conjugation systems, and enables the switch from the E1-E2 complex to the E2-E3 complex to allow conjugation reactions to proceed (*44, 45*). To confirm the binding of UFL1^UBS^ to UFC1 and compare its affinity with that of UBA5^UBS^, we performed isothermal titration calorimetry (ITC) between UFC1 and UFL1^UBS^. As indicated in Fig. 2G, the K_d_ value between UFL1^UBS^ and UFC1 was 0.41 μM, confirming that UFL1^UBS^ binds to UFC1, and its affinity was 2~2.5-fold higher than that of full-length UBA5 (*36*) and 6-fold higher than that of UBA5^UBS^ (*39*). These observations suggest that during ufmylation, UFC1 changes its binding partner from UBA5 to UFL1 to complete the conjugation reaction.

### Structural mechanism of substrate-specificity switching by CDK5RAP3

RPL26 is a major target of ufmylation, and the UFL1-UFBP1-CDK5RAP3 complex plays a critical role in this process (*13, 29, 30*). RPL26 ufmylation is enhanced by treatment with anisomycin, which prevents protein synthesis by inhibiting the peptidyl transferase center A site (*30, 46*). We confirmed that anisomycin treatment increased the levels of both di- and mono-ufmylated RPL26 (Fig. 3A). To investigate the effect of UFL1 mutants characterized by defective interaction with UFC1, UFBP1, or CDK5RAP3 during anisomycin-induced RPL26 ufmylation, we expressed each mutant in *UFL1* KO HEK293T cells. While ufmylation was completely suppressed by *UFL1* ablation, it was restored by expression of wild-type UFL1 (Fig. 3A). This restoration was not observed following the expression of any of the mutants (Fig. 3A), indicating that UFL1 interactions with UFC1, UFBP1, and CDK5RAP3 are required for the ufmylation of RPL26. UFBP1 is also ufmylated at Lys267, which enhances its ligase activity toward two other UFM1 substrates, ASC1 (*47*) and CYB5R3 (*27*). We noticed that UFL1^CDK5RAP mutant^, which showed defective interaction with CDK5RAP3, promoted the ufmylation of UFBP1 (Fig. 3A), which occurred even following anisomycin exposure (Fig. 3A). These results suggest that CDK5RAP3 may change the substrate specificity of the E3 enzyme. To study the role of CDK5RAP3 in substrate specificity, we predicted the structure of the UFL1-UFBP1-CDK5RAP3-UFC1 complex using AF2. In the absence of CDK5RAP3, UFBP1 Lys267 was exposed, and UFC1 bound to UFL1^UBS^ was identified near UFBP1 Lys267 (Fig. 3B). On the other hand, CDK5RAP3 binding dramatically changed the location of UFC1 so that it was distal to UFBP1 Lys267, by (1) narrowing the space around UFBP1 Lys267 via the T-dumbbell, and (2) binding to UFC1 using the IDR (Fig. 3C). An *in vitro* pull-down assay showed that recombinant CDK5RAP3, but not the UFC1-interacting defective mutants (CDK5RAP3^I267A W269A^, CDK5RAP3^UFC1 mutant^), bound to recombinant UFC1 (Fig. S5A). Additionally, while the immunoprecipitant prepared from *CDK5RAP3*-deficient cells expressing FLAG-tagged CDK5RAP3 contained endogenous UFC1, the level was much lower in the case of CDK5RAP3^UFC1 mutant^-expressing cells (Fig. S5B). These structural models, together with biochemical analyses, suggest that CDK5RAP3 impairs the ufmylation of UFBP1 Lys267 by positioning UFC1 away from the residue, and drives the ufmylation toward RPL26.

**Figure 3.**
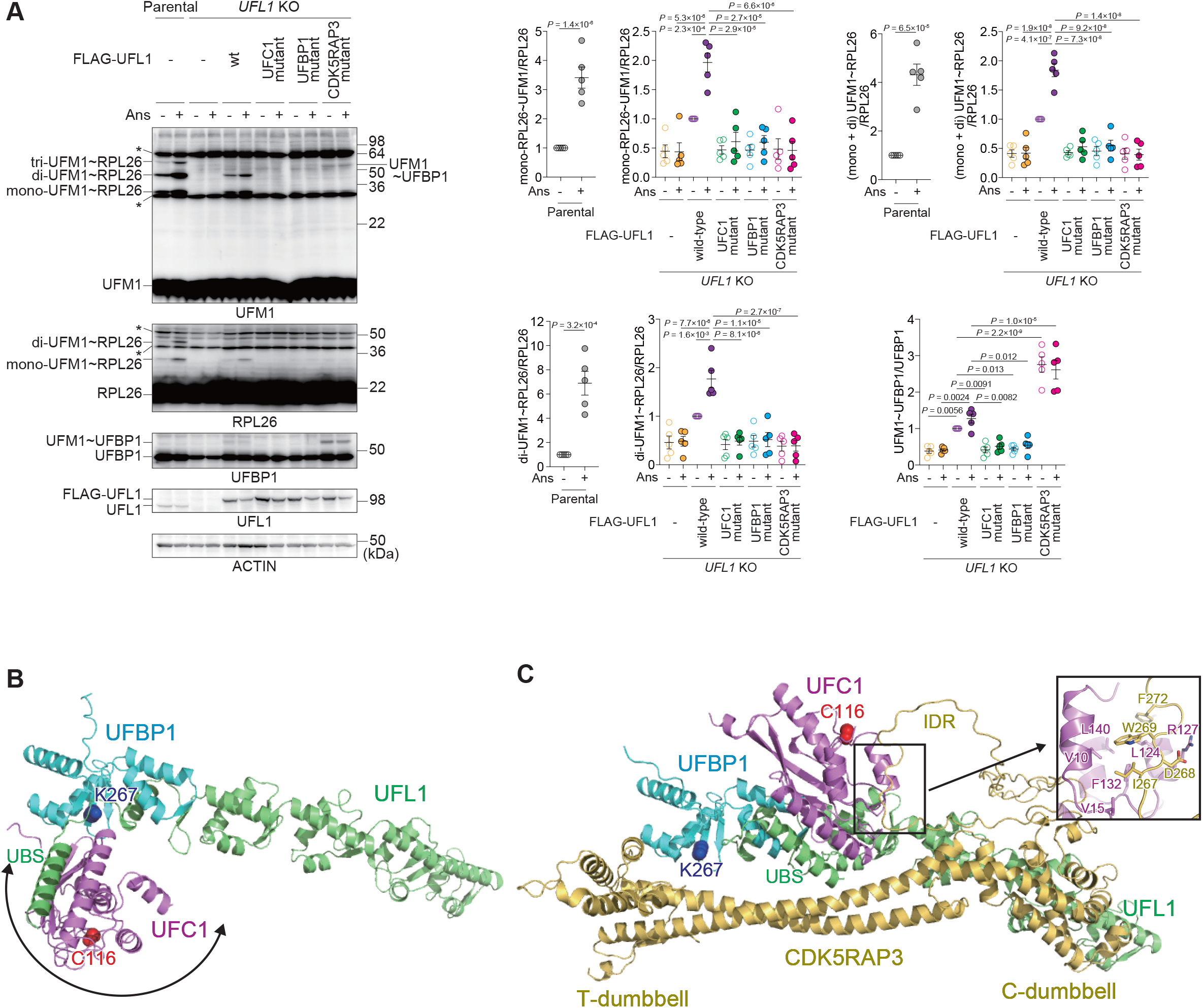
Substrate specificity switching by the UFM1 E3 component CDK5RAP3. **(A)** Immunoblot analysis. Wild-type UFL1 and UFC1-interaction-defective (UFL^UFC1 mutant^), UFBP1-interaction-defective (UFL^UFBP1 mutant^), and CDK5RAP3-interaction-defective UFL1 mutant (UFL^CDK5RAP3 mutant^) were transfected in *UFL1*-deficient HEK293T cells. Forty-eight hours after transfection, the cells were treated with 200 nM anisomycin (Ans) for 1 h and then lysed. The cell lysates were subjected to SDS-PAGE followed by immunoblot analysis with the indicated antibodies. Bar graphs show the results of quantitative densitometric analysis of ufmylated RPL26 (mono- and di-ufmylated RPL26) relative to free RPL26 (*n* = 3), and of ufmylated UFBP1 relative to free UFBP1 (*n* = 3). Data are means ± s.e. Statistical analysis was performed by Welch’s *t*-test. Data shown are representative of three separate experiments. **(B, C)** Predicted structures of the UFL1(1-302)-UFBP1(209-314)-UFC1 complex (B) and the UFL1(1-302)-UFBP1(209-314)-CDK5RAP3-UFC1 complex using AF2 (C). The positioning of UFC1 in (B) is variable due to the flexible nature of UFL1^UBS^. The side chains of UFBP1 Lys267 and UFC1 Cys116 are shown with a space-filling model. The inlet in (C) indicates the detailed interactions between the CDK5RAP3 IDR and UFC1.

### Interaction of CDK5RAP3 with UFM1

We recently showed that UFBP1 possesses a UFM1-interacting motif (UFIM) at the IDR N-terminal to the WH1 domain and binds to UFM1 in a manner similar to UBA5^UFIM^ (*27, 48*) (Fig. S6A and B). Unexpectedly, the E3 component CDK5RAP3, but not UFL1, was also predicted by AF2 to bind to UFM1 using the IDR inserted in the C-dumbbell (Fig. S6C). This binding process is similar to that of UFBP1^UFIM^ and UBA5^UFIM^, forming an intermolecular β-sheet with UFM1 β2 and inserting two hydrophobic residues into the hydrophobic pockets of UFM1 (Fig. S6D). Thus, this binding sequence in CDK5RAP3 was also named UFIM. An *in vitro* pull-down assay revealed that recombinant CDK5RAP3, but not the UFIM mutants (CDK5RAP3^I321A^, CDK5RAP3^UFIM mutant^), bound to recombinant UFM1 (Fig. S6E). To investigate the interaction of CDK5RAP3 with UFM1 in cells, we expressed wild-type CDK5RAP3 or CDK5RAP3^UFIM mutant^ in *CDK5RAP3*-deficient cells and then performed an immunoprecipitation assay. As shown in Fig. S6F, the immunoprecipitants from cells expressing wild-type CDK5RAP3, but not CDK5RAP3^UFIM mutant^, contained both free UFM1 and UFC1. We hypothesized that in addition to the interaction of CDK5RAP3 with UFC1 (Fig. 3C, Fig. S5), an intermediate consisting of a UFC1-UFM1 thioester binds to CDK5RAP3. To prove this, we utilized UFC1^C116S^, in which the active site cysteine is substituted with serine. Instead of the thioester bond, the UFC1^C116S^ formed an *O*-ester bond with UFM1, which was stable even under reducing conditions (*6*). When MYC-tagged UFC1^C116S^ and GFP-tagged UFM1 were co-expressed with wild-type or UFIM mutant CDK5RAP3 in *CDK5RAP3* KO cells, CDK5RAP3 interacted with GFP-UFM1-MYC-UFC1^C116S^ (Fig. S6G). The binding affinity was lower in the case of the CDK5RAP3^UFIM mutant^ (Fig. S6G). Very recently, *Arabidopsis thaliana* CDK5RAP3 was shown to interact with UFM1 directly using non-canonical LIRs (*49*). Since the UFIM sequence we identified in human CDK5RAP3 is not conserved in *A. thaliana*, the recognition mode of UFM1 by CDK5RAP3 may have diversified over the course of evolution.

CDK5RAP3 has the ability to interact with both UFC1 and UFM1-charged UFC1 (Figs. S5 and S6). These interactions may have a positive effect on RPL26 ufmylation. However, both UFC1- and UFM1-binding mutants of CDK5RAP3 promoted the formation of di- and tri-ufmylated RPL26 (data not shown), both of which are hardly observed under normal conditions. This suggests that binding of CDK5RAP3 to both UFC1 and UFM1-charged UFC1 may suppress di- and tri-ufmylation. Since di-ufmylated RPL26 has a high affinity to UFBP1 (refer to Fig. 4A), the binding of CDK5RAP3 to UFC1 and UFM1 negatively regulates the stable association of the UFM1 E3 complex with the 60S ribosomal subunit by inhibiting di-ufmylation of RPL26.

**Figure 4.**
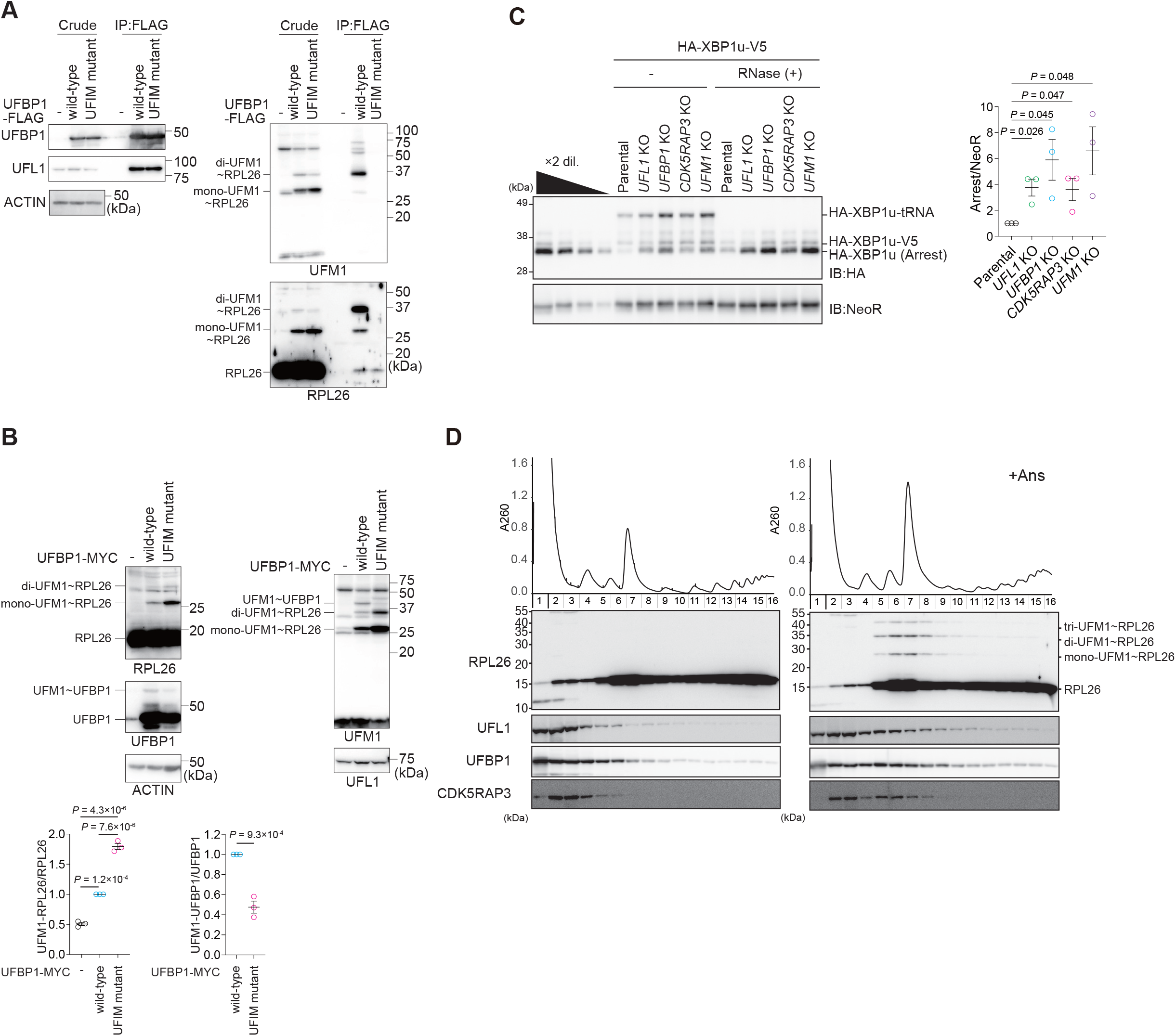
Anisomycin treatment induces association of the E3 complex with the 60S ribosomal subunit. **(A)** Immunoprecipitation assay. FLAG-tagged wild-type UFBP1 or UFM1-interaction-defective UFBP1 mutant (UFBP1^UFIM1 mutant^) was transfected into *UFBP1-* and *UFSP2*-deficient HEK293T cells. Forty-eight hours after transfection, cells were lysed and immunoprecipitated with anti-FLAG-M2 gel; the immunoprecipitants were then subjected to immunoblot analysis with the indicated antibodies. Data shown are representative of three separate experiments. **(B)** Immunoblot analysis. MYC-tagged wild-type UFBP1 or UFBP1^UFIM1 mutant^ was transfected into *UFBP1*-deficient HEK293T cells. Forty-eight hours after transfection, cells were lysed, and the cell lysates were subjected to SDS-PAGE followed by immunoblot analysis with the indicated antibodies. Bar graphs show the results of quantitative densitometric analysis of ufmylated RPL26 (mono- and di-ufmylated RPL26) relative to free RPL26 (*n* = 3) and of ufmylated UFBP1 relative to free UFBP1 (*n* = 3). Data are means ± s.e. Statistical analysis was performed by Welch’s *t*-test. Data shown are representative of three separate experiments. **(C)** The indicated KO cells were transfected with the *HA-XBP1u-V5* plasmid. Twenty-four hours after transfection, the cells were lysed. The cell lysates were subjected to neutral PAGE followed by immunoblot analysis with the indicated antibodies. (Left) The free peptide and peptidyl-tRNA (pep-tRNA) arrest products were detected by western blotting using antibodies against HA and NeoR. Samples treated with RNase to digest the tRNA moiety of pep-tRNA are indicated by (+). (Right) The relative levels of the HA-XBP1u arrest products with p values were determined by three independent experiments. **(D)** Cells were treated with 0.1 μg/ml anisomycin (Ans) for 30 min and then lysed. Ribosomes were separated by ultracentrifugation through sucrose density gradients. Protein samples were prepared from gradient fractions and analyzed by western blotting using antibodies against RPL26, UFM1, UFL1, UFBP1, and CDK5RAP3.

### Association of the E3 UFM1 complex with the 60S ribosomal subunit

Next, to test the interaction of UFBP1 with UFM1 in cells, we generated *UFBP1 UFSP2* double knockout (DKO) HEK293T cells (Fig. S3B) and used them to express wild-type UFBP1 and UFIM mutants (UFBP1^F196A V198A^, UFBP1^UFIM mutant^). Both proteins were expressed at a similar level (Fig. 4A). In cells lacking *UFSP2*, the ufmylated proteins accumulated (*16*). Indeed, when wild-type UFBP1 was expressed in the *UFBP1 UFSP2* DKO HEK293T cells, we observed ufmylated RPL26 even without exposure to anisomycin (Fig. 4A). An immunoprecipitation assay showed that wild-type UFBP1, but not UFBP1^UFIM mutant^, interacted with di- and mono-ufmylated RPL26, in particular di-ufmylated RPL26 (Fig. 4A). We therefore sought to determine the effect of the UFBP1 UFIM mutant on ufmylation of RPL26 and UFBP1. When UFBP1^UFIM mutant^ was overexpressed in *UFBP1*-deficient cells, UFBP1 ufmylation was impaired, and this enhanced the ufmylation of RPL26 (Fig. 4B). This implies that the stable association of UFBP1 with ufmylated RPL26 may trap the UFM1 E3 on the ribosome and then suppress RPL26 ufmylation.

What is the significance of the interaction between UFBP1 and ufmylated RPL26? Anisomycin treatment increases the ZNF598-mediated ubiquitination of uS10 and eS10 and their disome formation (*50–53*) and also increases ufmylation, suggesting that ufmylation is required for ER-RQC. We examined the role of the UFL1-UFBP1-CDK5RAP3 E3 complex in RQC-mediated degradation of nascent polypeptides derived from stalled ribosomes. Ribosome stalling during the translation of *XBP1u* mRNA contributes to the efficient targeting of *XBP1u* mRNA to the ER membrane (*54, 55*). Ribosome stalling at the *XBP1u* stalling sequence induces ER-RQC, and nascent polypeptides derived from stalled ribosomes are degraded in an LTN1-dependent manner (*56*). To evaluate the role of the UFL1-UFBP1-CDK5RAP3 E3 complex for ER-RQC, we investigated LTN1-mediated degradation of the nascent XBP1u polypeptide in UFM1 E3 KO cells. The levels of nascent XBP1u polypeptide were increased in *UFL1, UFBP1, CDK5RAP3*, and *UFM1* KO cells (Fig. 4C), indicating that ufmylation is required for ER-RQC.

We next asked whether the association of the UFL1-UFBP1-CDK5RAP3 E3 complex with the 60S ribosomal subunit is facilitated by anisomycin treatment, which increases both ribosome collisions (*57*) and the levels of di- and mono-ufmylated RPL26 (Fig. 3A). To verify the interaction of the subunits of the UFM1 E3 complex with ribosomes, we performed polysome analysis. The association of E3 components with the 60S subunit was estimated by western blotting using the fractions obtained after centrifugation in sucrose gradients. Without anisomycin treatment, UFL1 and CDK5RAP3 were distributed mainly in the ribosome-free fraction whereas UFBP1 was found in both the ribosome-free and 60S fractions. Upon anisomycin treatment, the distribution of these factors in the 60S fraction was significantly increased (Fig. 4D). Given that the interaction of UFL1 with CDK5RAP3 was observed even without anisomycin treatment (Figs. 1 and 2), and UFL1 formed a stable complex with the ER-localizing protein UFBP1 (*13, 28*), we propose that the E3 UFM1 complex associates with the 60S subunit and that this association is facilitated by anisomycin treatment.

### The interaction of UFBP1 with ufmylated RPL26 is crucial for ER-RQC

We next investigated which component of the UFM1 E3 complex is indispensable for the anisomycin-induced association of E3 complex with the 60S ribosome. Polysome analysis revealed that deletion of one of the three E3 components disrupted the association of the other components with the 60S subunit (Fig. 5A-D). In *UFL1* KO cells, the expression level of CDK5RAP3 was significantly reduced, as was the association of UFBP1 with the 60S subunit (Fig. 5B). In *UFBP1* KO cells, neither UFL1 nor CDK5RAP3 was associated with the 60S subunit (Fig. 5C). In *CDK5RAP3* KO cells, the association of UFBP1 and UFL1 with the 60S subunit was reduced (Fig. 5D), suggesting that ufmylation by the E3 complex facilitates its interaction with the 60S subunit. Since UFBP1 associates with ufmylated RPL26 via UFBP1^UFIM^-UFM1 interaction, we next examined whether the defect of the interaction affects the association of the UFM1 E3 complex with the 60S subunit. Compared with wild-type UFBP1 expression, the association of UFL1 and CDK5RAP3 with the 60S subunit was significantly reduced in *UFBP1*-deficient cells expressing UFBP1^UFIM mutant^ (Fig. 6A), indicating that the interaction of UFBP1 with ufmylated RPL26 via the UFIM contributes to the association of the E3 complex with the 60S subunit. We evaluated the role of UFBP1^UFIM^ for ER-RQC and found that the levels of nascent XBP1u polypeptides were increased in *UFBP1*-deficient cells expressing UFBP1^UFIM mutant^ (Fig. 6B), indicating that the interaction of UFBP1 with ufmylated RPL26 is required for ER-RQC.

**Figure 5.**
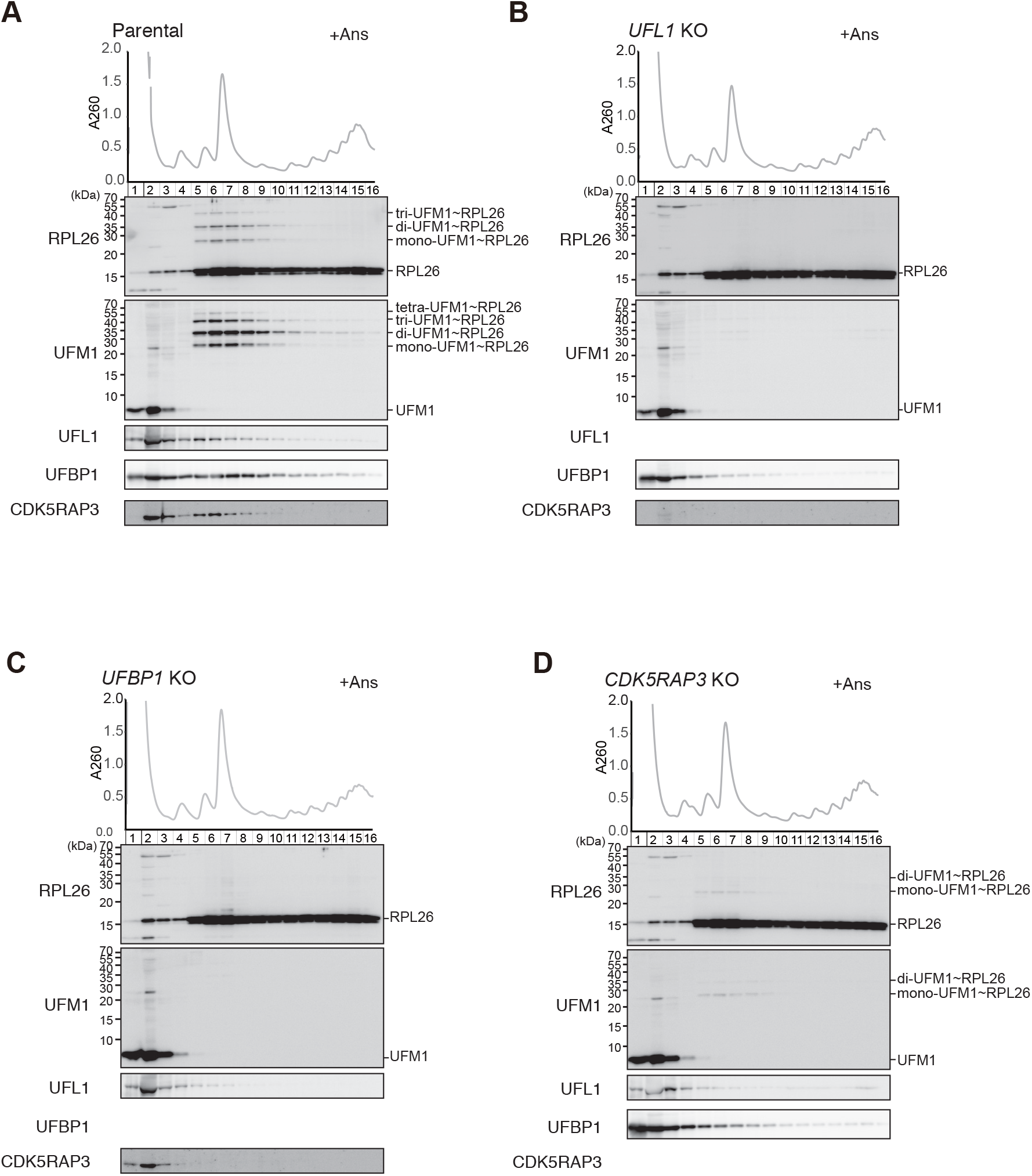
UFBP1 is indispensable for the association of E3 components with the 60S ribosomal subunit upon anisomycin treatment. Parental HEK293T cells **(A)** or *UFL1-* **(B)**, *UFBP1-* **(C)**, and *CDK5RAP3*- **(D)** KO cells were treated with 0.1 μg/ml anisomycin (Ans) for 30 min, and then lysed. Ribosomes were separated by ultracentrifugation through sucrose density gradients. Protein samples were prepared from gradient fractions and analyzed by western blotting using antibodies against RPL26, UFM1, UFL1, UFBP1, and CDK5RAP3.

**Figure 6.**
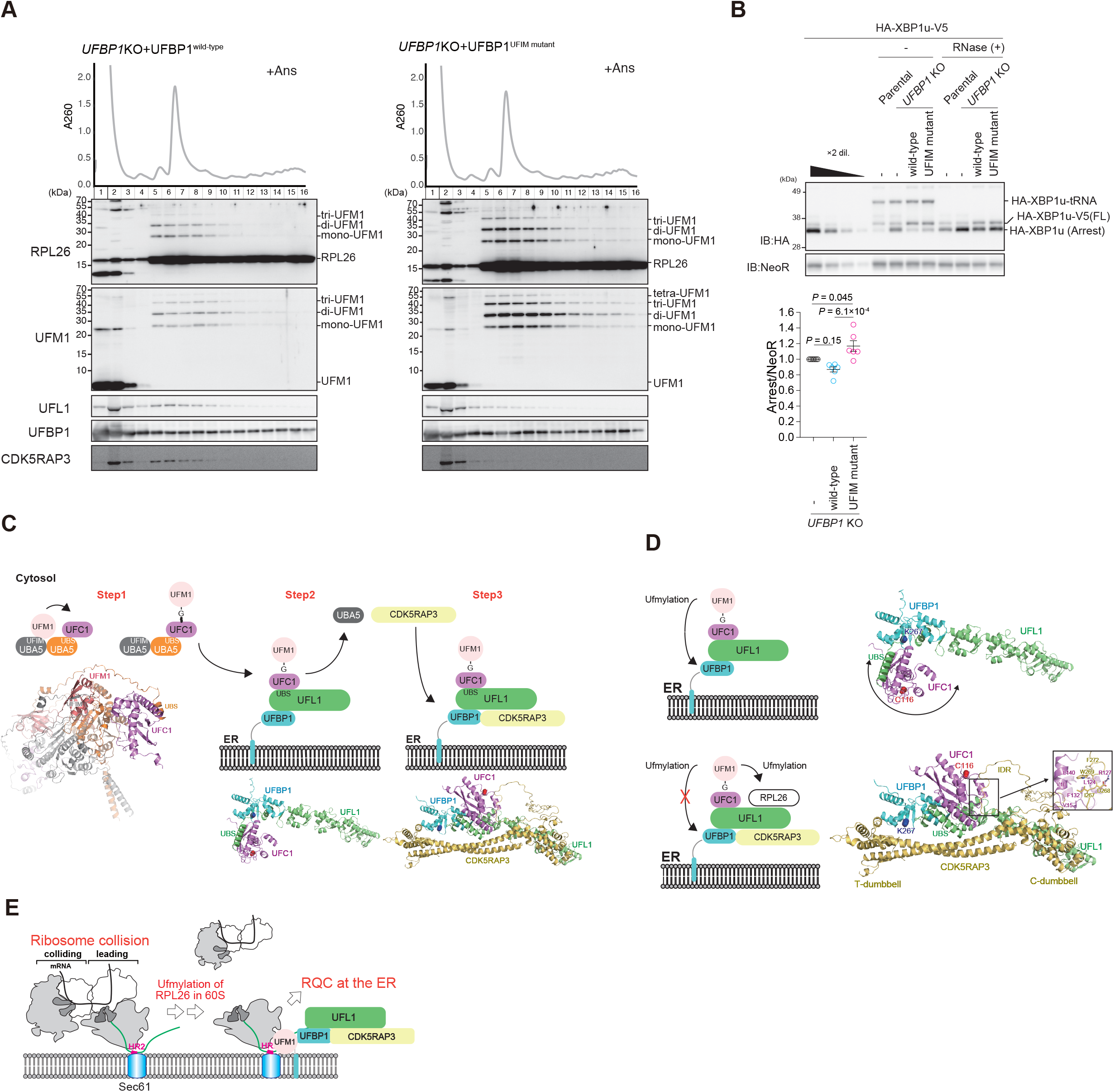
The interaction of UFBP1 with ufmylated RPL26 is indispensable for ER-RQC. **(A)** *UFBP1* KO cells were transfected with plasmids expressing wild-type UFBP1 or UFM1-interaction-defective UFBP1. Twenty-four hours after transfection, the cells were treated with 0.1 μg/ml anisomycin (Ans) for 30 min and then lysed. Ribosomes were separated by ultracentrifugation through sucrose density gradients. Protein samples were prepared from gradient fractions and analyzed by western blotting using antibodies against RPL26, UFM1, UFL1, UFBP1, and CDK5RAP3. **(B)** Wild-type UFBP1 or UFBP1^UFIM mutant^ was co-transfected with the *HA-XBP1u-V5* plasmid in *UFBP1* KO cells. Twenty-four hours after transfection, the cells were lysed. (Left) The cell lysates were subjected to neutral PAGE followed by immunoblot analysis with the indicated antibodies. The free peptide and the peptidyl-tRNA (pep-tRNA) arrest products were detected by western blotting using an anti-HA antibody. Samples treated with RNase to digest the tRNA moiety of pep-tRNA are indicated by (+). Essentially the same results were obtained in two independent experiments. (Right) The relative levels of the HA-XBP1u arrest products with p values were determined by three independent experiments. (**C**) Schematic model of the enzymatic cascades during RPL26 ufmylation. Each protomer within the UBA5 homodimer binds UFM1 and UFC1 via UFIM and UBS, respectively, and catalyzes the formation of the UFM1-UFC1 thioester intermediate. UFL1-UFBP1 anchored to the ER recruits the UFM1-UFC1 intermediate via UFL1^UBS^ binding to UFC1. Finally, CDK5RAP3 is recruited to the E3 complex to complete the organization of the pentameric E3 complex on the ER. (**D**) Schematic model of substrate switching of the UFM1 E3 ligase. In the absence of CDK5RAP3, the UFM1-UFC1 intermediate can freely change its position and thus access and ufmylate UFBP1 K267. Upon CDK5RAP3 binding, the position of the UFM1-UFC1 intermediate is locked away from UFBP1 K267, preventing its ufmylation and promoting RPL26 ufmylation instead. (**E**) Schematic model of ER-RQC through the UFM1 E3 ligase. Upon the treatment with anisomycin, an inducer of disome formation (*i.e*., ribosomal collision), the UFM1 E3 complex stably associated with ufmylated RPL26 on the 60S ribosomal subunit through the UFM1-interacting region of UFBP1, and the association was required for ER-RQC.

## Discussion

Based on multiple structural analyses of the ternary UFM1 E3 complex, E2, and UFM1 using AF2, as well as biochemical analyses, we propose a sequential organization model of the E3 complex on the ER as follows. A complex consisting of UBA5 and UFC1, which are the E1 and E2 enzymes associated with UFM1, respectively, initially generates a UFM1-UFC1 thioester intermediate. Next, a complex composed of the E3 components UFL1 and UFBP1 receives the UFM1-UFC1 intermediate from UBA5 at the ER by competitive binding of UFL1^UBS^ to UFC1. Finally, the UFL1-UFBP1 complex with the UFM1-UFC1 intermediate recruits CDK5RAP3 to create the UFL1-UFBP1-CDK5RAP3-UFM1-UFC1 pentameric complex on the ER, which relocates the UFM1-UFC1 intermediate away from UFBP1 Lys267 to promote ufmylation of RPL26 (Fig. 6C). On the other hand, if CDK5RAP3 is unavailable to bind to the UFL1-UFBP1 complex, the UFM1-UFC1 intermediate can freely change its position relative to the E3 complex to ufmylate UFBP1 Lys267. This model suggests that CDK5RAP3 functions as an accessory component of the E3 complex and switches the E3 complex specificity from UFBP1 to RPL26 (Fig. 6D).

Anisomycin treatment promotes RPL26 ufmylation and organization of the pentameric E3 complex on the ER. How is this E3 complex organization regulated by anisomycin? Since the UFL1-UFBP1, UFC1-UFL1, and UFL1-CDK5RAP3 interactions were observed irrespective of anisomycin treatment (Figs. 1 and 2), and the UFL1-UFBP1 complex was always anchored to the ER, recruitment of the UBA5-UFC1 complex and CDK5RAP3 to the ER should be enhanced in response to anisomycin. UBA5 has been identified as a GABARAP-interacting protein, and a GABARAP family protein, GABARAPL2 translocates UBA5 to the ER through the interaction with ACSL3, an ER-associated protein and lipid droplet biogenesis factor (*6, 58, 59*). If the anchoring of UBA5 at the ER is impaired, the UBA5 protein level decreases, implying that ACSL3 serves as a stabilizing factor for GABARAP-binding proteins (*58*). Since CDK5RAP3 also forms a complex with LC3-family proteins through non-canonical LIRs (*32*), it may localize on the ER in a manner similar to UBA5 and is then transferred to UFL1. This hypothesis is supported by the finding in this study that free CDK5RAP3, which does not localize on the ER, became unstable, which was also the case with UBA5 (Fig. 1D and F). If concomitant recruitment of UBA5 and CDK5RAP3 to the ER is promoted by anisomysin treatment, this should increase RPL26 ufmylation. While RPL26 ufmylation is involved in ER-RQC and/or nascent peptide quality control on the ER (*29–31*), ufmylation of UFBP1 contributes to ER-phagy (*27*). When the threshold level of ER-RQC is exceeded, ER-phagy may induce the removal of ER instances that contain a number of stalled ribosomes. To accomplish this, UFM1 E3 may have a substrate-switching mechanism involving suppression of the ER localization of CDK5RAP3, but not UBA5. CDK5RAP3 was reported to have three non-canonical LIRs (IDWG sequences), one of which is an exact match for the UFC1 binding region (*32*) (Fig. 3C). Therefore, sufficient binding of GABARAP to the LIRs would competitively inhibit the interaction of CDK5RAP3 with UFC1, thereby promoting dissociation of CDK5RAP3 from the UFC1-charged E3 complex. Further analysis is needed to clarify these regulatory mechanisms.

Another important finding of this study is that the UFL1-UFBP1-CDK5RAP3 E3 complex stably binds to ufmylated RPL26 via UFBP1^UFIM^-UFM1 interaction, which is an additional function of the E3 complex separate from its ligase activity. What, then, is the significance of the stable association of UFBP1 with the 60S ribosomal subunit? Even though LTN1-mediated degradation of XBP1u stalled products was impaired in *UFBP1*-deficient cells expressing UFBP1^UFIM mutant^ (Fig. 6B), the ufmylation of RPL26 in the 60S subunit was promoted (Fig. 6A). These results indicate that the stable association of the E3 complex with the 60S subunit via UFBP1 binding to UFM1-RPL26, but not the ligase activity of this complex, is crucial for ER-RQC (Fig. 6E). The mechanism by which the LTN1-mediated degradation of the polypeptide on the 60S subunit depends on the interaction of UFBP1 with this subunit remains unknown. Given that RPL26 is located close to the translocon (*60*), the recruitment of NEMF and/or LTN1 to the 60S subunit may require the interaction of UFBP1 with the 60S subunit through ufmylated RPL26. The ufmylation of the 60S subunit containing peptidyl-tRNA, an LTN1 substrate, leads to the interaction of UFBP1 with ufmylated RPL26 through the UFIM, resulting in a stable association of the UFM1 E3 complex with the 60S subunit and the binding of LTN1 for the ubiquitination of the peptidyl-tRNA on this subunit. Structural analysis of the 60S-UFM1 E3 complex will provide fundamental information to clarify the crucial role of UFM E3 in quality control that eliminates aberrant products derived from ribosome collision.

## Materials and methods

### Structural prediction using AF2 with the AlphaFold-Multimer mode

Complex structures were predicted using AlphaFold2 v2.2.0 (downloaded on May 16, 2022 from https://github.com/deepmind/alphafold) installed on a local computer (Sunway Technology Co., Ltd.) (*61*). The predictions were run using the AlphaFold-Multimer mode (*42*), with five models and a single seed per model, and default multiple sequence alignment generation using the MMSeqs2 server (*62*). The unrelaxed predicted models were subjected to an Amber relaxation procedure and the relaxed model with the highest confidence based on predicted LDDT scores was selected as the best model and used for figure preparation (*61*). Structural figures were prepared using PyMOL (http://www.pymol.org/pymol).

### Cell culture

HEK293T cells (ATCC CRL-3216) were grown in Dulbecco’s modified Eagle medium containing 10% fetal bovine serum, 2 mM L-glutamine, 5 U/ml penicillin, and 50 μg/ml streptomycin. To introduce expression vectors, HEK293T cells were transfected with PEI MAX (Polysciences, Inc., Warrington, PA, USA). *UFL1* (5’-CCAGCGGGCGCAGTTCGCCG-3’) or *UFBP1* (5’-GTAGCGGCGGCTCTGCTAGT’) guide RNA was designed using the CRISPR Design tool (http://crispr.mit.edu/) and subcloned into pX330-U6-Chimeric_BB-CBh-hSpCas9 (Addgene #42230), a human codon-optimized SpCas9 and chimeric guide RNA expression plasmid. To generate *UFL1, UFC1, CDK5RAP3*. and *UFBP1 UFSP2* KO HEK293T cells, HEK293T or *UFSP2* KO HEK293T cells (*16*) were transfected with the aforementioned pX330 vectors together with pEGFP-C1 (#6084-1, Clontech Laboratories, Mountain View, CA, USA) and cultured for 2 d. GFP-positive cells were sorted and expanded. Ablation of *UFL1, UFC1*, or *CDK5RAP3* was confirmed by a heteroduplex mobility assay followed by immunoblot analysis with anti-UFL1, anti-UFC1, or anti-CDK5RAP3 antibody. *UFC1- (23), UFBP1-* (*27*), *CDK5RAP3-* (*27*), and *UFSP2*-deficient HEK293T cells (*16*) were used in this study. HEK293T and HeLa cells were authenticated by STR profile. All cell lines were tested for mycoplasma contamination.

### Immunoblot and immunoprecipitation analysis

Cells were lysed with ice-cold TNE buffer containing 1% NP40, 1% TX-100, and protease inhibitors. The lysates were centrifuged at 20,000 *g* for 10 min at 4°C, and the resulting supernatants were used as samples for immunoblot analysis. Samples were subjected to SDS-PAGE, then transferred to a polyvinylidene difluoride (PVDF) membrane (IPVH00010; Merck Millipore, Burlington, MA, USA). Antibodies against UFM1 (ab109305, Abcam, Cambridge, UK; 1:1000), UFL1 (A303-456A; Bethyl Laboratories, Montgomery, TX, USA; 1:1000), UFBP1 (21445-1-AP, Proteintech; 1:1000), CDK5RAP (H00080279-M01; Novus Biologicals, Englewood, CO, USA; 1:500), UFSP2 (ab185965, Abcam; 1:1000), PRL26 (ab59567, Abcam; 1:1000), ACTIN (A1978; Sigma-Aldrich, Burlington, MO, USA; 1:2000), and FLAG (M185-3L, Medical & Biological Laboratories, Tokyo, Japan; 1:1000) were purchased from the indicated suppliers. Blots were incubated with horseradish peroxidase-conjugated goat anti-mouse IgG (H+L) (115-035-166, Jackson ImmunoResearch Laboratories, Inc., West Grove, PA, USA; 1:10000) or goat anti-rabbit IgG (H+L) (111-035-144, Jackson ImmunoResearch Laboratories, Inc.; 1:10000), and visualized by chemiluminescence. Band density was measured using the software Multi Gauge V3.2 (FUJIFILM Corporation, Tokyo, Japan). For immunoprecipitation analysis, cells were lysed in 300 μl of Immunoprecipitation (IP) buffer (20 mM Tris-HCl [pH 7.5], 150 mM NaCl, 1 mM EDTA, 1% NP40, and 1% TX-100) containing Protease Inhibitor Cocktail (Roche, Basel, Switzerland), and the lysates were then centrifuged at 20,000 *g* for 10 min at 4°C to remove debris. In the next step, 200 μl of IP buffer and 10 μl of anti-FLAG M2 Affinity Agarose Gel (A2220, Merck Millipore) were added to the 200 μl of lysate, and the mixture was mixed under constant rotation for 3 h at 4°C. The immunoprecipitates were washed five times with ice-cold IP buffer. The complex was boiled for 5 min in SDS sample buffer in the presence of ß-mercaptoethanol to elute proteins.

### Isothermal titration calorimetry (ITC)

For ITC experiments, UFC1 prepared for pull-down assay and synthesized UFL1 (^1^MADAWEEIRRLAADFQRAQFA^21^, from Toray Research Center, Tokyo, Japan) were subjected to size-exclusion chromatography with 20 mM HEPES (pH 6.8) and 150 mM sodium chloride using a Superdex 200 column (Cytiva, Tokyo, Japan). ITC experiments were performed using a Microcal iTC200 calorimeter (Malvern Panalytical, Malvern, Worcestershire, UK), with stirring at 750 rpm at 25°C. The cell and syringe were filled with 30 μM UFC1 and 300 μM UFL1(1-21), respectively. The titration involved 18 injections of 2 μl of the syringe sample at intervals of 120 s into a cell after one injection of 0.4 μl of syringe sample. MicroCal Origin 7.0 software was used for data analysis. Thermal measurement data of the first syringe sample injections were removed from the analysis. Thermal titration data were fit to a single-site binding model, which determines thermodynamic parameter the enthalpy (ΔH), dissociation constant (Kd), and stoichiometry of binding (N). The error of each parameter shows the fitting error.

### Pull-down assay

Recombinant GST-fused UFM1, UFC1, UFBP1, UFBP1^F196A V198A^, CDK5RAP3, CDK5RAP3^I321A^, and CDK5RAP3^I267A W269A^ were produced in *Escherichia coli* and purified by chromatography on glutathione-Sepharose 4B resin (Amersham Biosciences, Arlington Heights, IL, USA). GST-tag was cleaved by precision protease (Amersham Biosciences). The purified proteins were mixed in TNE buffer (50 mM Tris-HCl [pH 7.5], 150 mM NaCl, 1 mM EDTA) containing 1% NP40 for 1 h at 4°C and then precipitated with glutathione Sepharose. The mixtures were washed five times with ice-cold TNE. The bound proteins were analysed by SDS-PAGE followed by Coomassie brilliant blue staining or immunoblot analysis.

### RQC assay using *XBP1u* staller sequence

RQC activity was monitored by the detection of the arrest product derived from a reporter *HA-XBP1u-V5* plasmid (*56*). The UFL1, UFBP1, CDK5RAP3, and UFM1 knockout cells were transfected with *HA-XBP1u-V5* plasmid, and cell lysates were prepared after 24 h. Protein samples of cell lysates with or without RNase treatment were separated by neutral PAGE followed by western blotting to detect the HA-XBP1u arrest products using the anti-HA antibody to identify peptidyl-tRNAs (*52, 56*). The levels of the HA-XBP1u-arrest products and the control NeoR proteins were quantified using dilution series. The relative levels of the HA-XBP1u arrest products with p values were calculated by three independent experiments.

### Sucrose density gradient ultracentrifugation and western blotting

Cells were treated with or without anisomycin at the indicated concentrations for 30 min (37°C, 5% CO_2_). Cells were washed with phosphate-buffered saline twice and lysed with lysis buffer (50 mM Tris–HCl [pH 6.8], 100 mM NaCl, 10 mM MgCl□, 1% NP-40, 2 mM 2-mercaptoethanol, 1 mM PMSF) supplemented with protease inhibitors (Thermo Fisher Scientific, Waltham, MA, USA) and centrifuged at 1,500 *g* for 5 min at 4°C. The HEK293 cell extracts were layered on top of the sucrose gradients and centrifuged at 201,000 *g* in a SW40 rotor for 2□h at 4°C. The gradients were then fractionated with a BioComp Piston Gradient Fractionator (BioComp Instruments, Fredericton, New Brunswick, Canada). The polysome profiles were generated by continuous absorbance measurement at 26□nm using a single-path UV-1 optical unit (ATTO Biomini UV-monitor, ATTO, Tokyo, Japan) connected to a chart recorder (ATTO digital mini-recorder). For the western blots, 500 μl of each fraction was mixed with 55.6 μL of 100% TCA and incubated for 20 min at 4°C. After centrifugation (15,000 rpm, 15 min, 4°C), the supernatant was removed and the pellet was washed with acetone and dissolved in 30 μl of 2× SDS sample buffer (100 mM Tris [pH 6.8], 4% w/v SDS, 20% glycerol, 0.02% BPB, and 50 mM DTT). Protein samples were separated by SDS-PAGE and transferred to a PVDF membrane (#IPVH00010, Millipore).

For neutral PAGE, cells were lysed with lysis buffer and centrifuged at 20,000 *g* for 10 min. Supernatants were collected, and equal amounts of total proteins were used as protein samples. For RNase(+) samples, RNase A (QIAGEN, Hilden, Germany) was added at a final concentration of 0.05 mg/ml and incubated on ice for 30 min. For RNase(-) samples, Milli-Q water was added instead. After incubation, 1x Sample Buffer (50 mM Tris [pH 6.8], 2% w/v SDS, 10% glycerol, 0.01% BPB, and 50 mM DTT) was added and heated for 5 min at 70°C. Proteins were separated by 15% PAGE under neutral pH conditions (pH 6.8) for 5 h with a 150 V constant voltage in MES-SDS buffer (50 mM MES, 50 mM Tris base, 3.465 mM SDS, and 1 mM EDTA), and were transferred to a PVDF membrane (#IPVH00010, Millipore).

### Statistical analysis

Values, including those displayed in the graphs, are presented as means ± s.e. Statistical analysis was performed using the unpaired t-test (Welch test) or Šidák’s multiple comparison test with GraphPad Prism ver 9.2.0 software (GraphPad software, Boston, MA, USA). A P value less than 0.05 was considered to indicate statistical significance.

## Acknowledgements

R.I. is supported by a Grant-in-Aid for Scientific Research (C) (22K06931). S.K-H. is supported by a Grant-in-Aid for Encouragement of Scientists (21H04163). T.I. is supported by AMED (JP20gm1110010, JP223fa627001), MEXT/JSPS KAKENHI (JP19H05281, 21H05277, 22H00401), and the Takeda Science Foundation. N.N.N. is supported by MEXT/JSPS KAKENHI (JP19H05707), JST CREST (JPMJCR20E3), and the Takeda Science Foundation. M.K. is supported by MEXT/JSPS KAKENHI (19H05706, 21H004771), AMED-CREST (22gm1410004h0003), and the Takeda Science Foundation. We would like to thank H. Morishita, R. Kurusu, and Y. Ichimura for critically reading this manuscript.

## Author Contributions

T.I., N.N.N., and M.K. designed and directed the study. R.I., S.I., and M.G. carried out the biochemical experiments. S.K-H. generated knockout cell lines. N.N.N conducted structural analyses with AlphaFold2. T.I., N.N.N., and M.K. wrote the manuscript. All authors discussed the results and commented on the manuscript.

---

**Supplementary Figure S1 Predicted structure of each UFM1 E3 component.**

Predicted three-dimensional structures of UFL1 (UniPlot O94874), UFBP1 (UniPlot Q96HY6), and CDK5RAP3 (UniPlot Q96JB5) downloaded from the AlphaFold Protein Structure Database. AF2 produces a per-residue confidence score (pLDDT) between 0 and 100, which is indicated by color (the color-score relationship is shown in the box; pLDDT scores between 0 and 50 are colored red).

**Supplementary Figure S2 Predicted structure of the UFM1 E3 complex.**

**(A)** Predicted three-dimensional structures of the UFL1(1-302)-UFBP1(209-314)-CDK5RAP3 complex by AF2. The top five models are shown superimposed. **(B)** Binding mode of the T-dumbbell of CDK5RAP3 to the N-terminal helix of UFL1 and the WH1 domain of UFBP1. The side chains involved in this interaction are shown with a stick model, where oxygen and nitrogen atoms are colored red and blue, respectively. Broken lines indicate possible electrostatic interactions.

**Supplementary Figure S3 Generation of knockout cell lines.**

**(A, B)** Immunoblot analysis. Generation of *UFL1* knockout (A) and *UFBP1 UFSP2* double knockout (B) cell lines. The indicated genotype cell lines were lysed, then subjected to SDS-PAGE followed by immunoblot analysis with the indicated antibodies. Data shown are representative of three separate experiments.

**Supplementary Figure S4 Predicted binding mode of UFM1 E2 UFC1 to E3 subunits.**

**(A)** Top-scored model of the full-length UFL1-UFC1 binary complex predicted by AF2. **(B)** Top five prediction models of the full-length UFL1-UFC1 binary complex are shown superimposed. **(C)** Structural prediction of the UFL1^UBS^-UFC1 binary complex. Top five models are shown superimposed. **(D)** Top-scored model of the UFBP1-UFC1 binary complex predicted by AF2. The right panel shows detailed interactions, where the side chains involved in the interaction are indicated with a stick model. Oxygen and nitrogen atoms are colored red and blue, respectively. Broken lines indicate possible electrostatic interactions. **(E)** Top five models of the UFBP1-UFC1 binary complex are shown superimposed. **(F)** Structural prediction of the UFL1-UFBP1-UFC1 ternary complex. The top five models are shown superimposed. **(G)** Comparison of the predicted structure of UFC1 (AF2) with the crystal structure (2Z6O) (left). Comparison of the predicted structure of free UFC1 and UFL1^UBS^-bound UFC1 (right).

**Supplementary Figure S5 CDK5RAP3 interacts with UFC1.**

**(A)** Pull-down assay. GST, GST-CDK5RAP3, or GST-CDK5RAP3^I267A W269A^ (CDK5RAP3^UFC1 mutant^) was immobilized on glutathione Sepharose. The Sepharose and recombinant UFM1 were incubated, then the supernatant and pulled-down products were subjected to SDS-PAGE followed by immunoblotting with anti-UFC1 antibody. Data shown are representative of three separate experiments. **(B)** Immunoprecipitation assay. FLAG-tagged wild-type CDK5RAP3 or CDK5RAP3^UFC1 mutant^ was transfected into *CDK5RAP3*-deficient HEK293T cells. Forty-eight hours after transfection, cells were lysed and immunoprecipitated with anti-FLAG-M2 gel; the immunoprecipitants were then subjected to immunoblot analysis with the indicated antibodies. Data shown are representative of three separate experiments.

**Supplementary Figure S6 CDK5RAP3 interacts with UFM1-charged UFC1.**

**(A)** Crystal structure of UFBP1^UFIM^ bound to UFM1 (PDB 7W3N). **(B)** Crystal structure of UBA5^UFIM^ bound to UFM1 (PDB 5HKH). **(C)** Structural prediction of the CDK5RAP3-UFM1 binary complex. The top five models are shown superimposed. **(D)** Close-up view of the interaction between CDK5RAP3^UFIM^ and UFM1 in the top-scored model. **(E)** Pull-down assay. GST, GST-CDK5RAP3, or GST-CDK5RAP3 UFIM mutant (CDK5RAP3^UFIM1 mutant^) was immobilized on glutathione Sepharose. The Sepharose and recombinant UFM1 were incubated, then the supernatant and pulled-down products were subjected to SDS-PAGE followed by immunoblotting with anti-UFM1 antibody. Data shown are representative of three separate experiments. **(F-G)** Immunoprecipitation assay. FLAG-tagged wild-type CDK5RAP3 or CDK5RAP3^UFIM1 mutant^ was transfected into *CDK5RAP3*-deficient HEK293T cells (F). FLAG-tagged wild-type CDK5RAP3 or CDK5RAP3^UFIM1 mutant^, together with MYC-tagged UFC1^C116S^ and GFP-tagged UFM1, were transfected into *CDK5RAP3*-deficient HEK293T cells (G). Forty-eight hours after transfection, cells were lysed and immunoprecipitated with anti-FLAG-M2 gel; the immunoprecipitants were then subjected to immunoblot analysis with the indicated antibodies. Data shown are representative of three separate experiments.

