## Supplementary Figures for "Mechanistic insights into the UFM1 E3 ligase complex in ufmylation and ribosome-associated protein quality control"

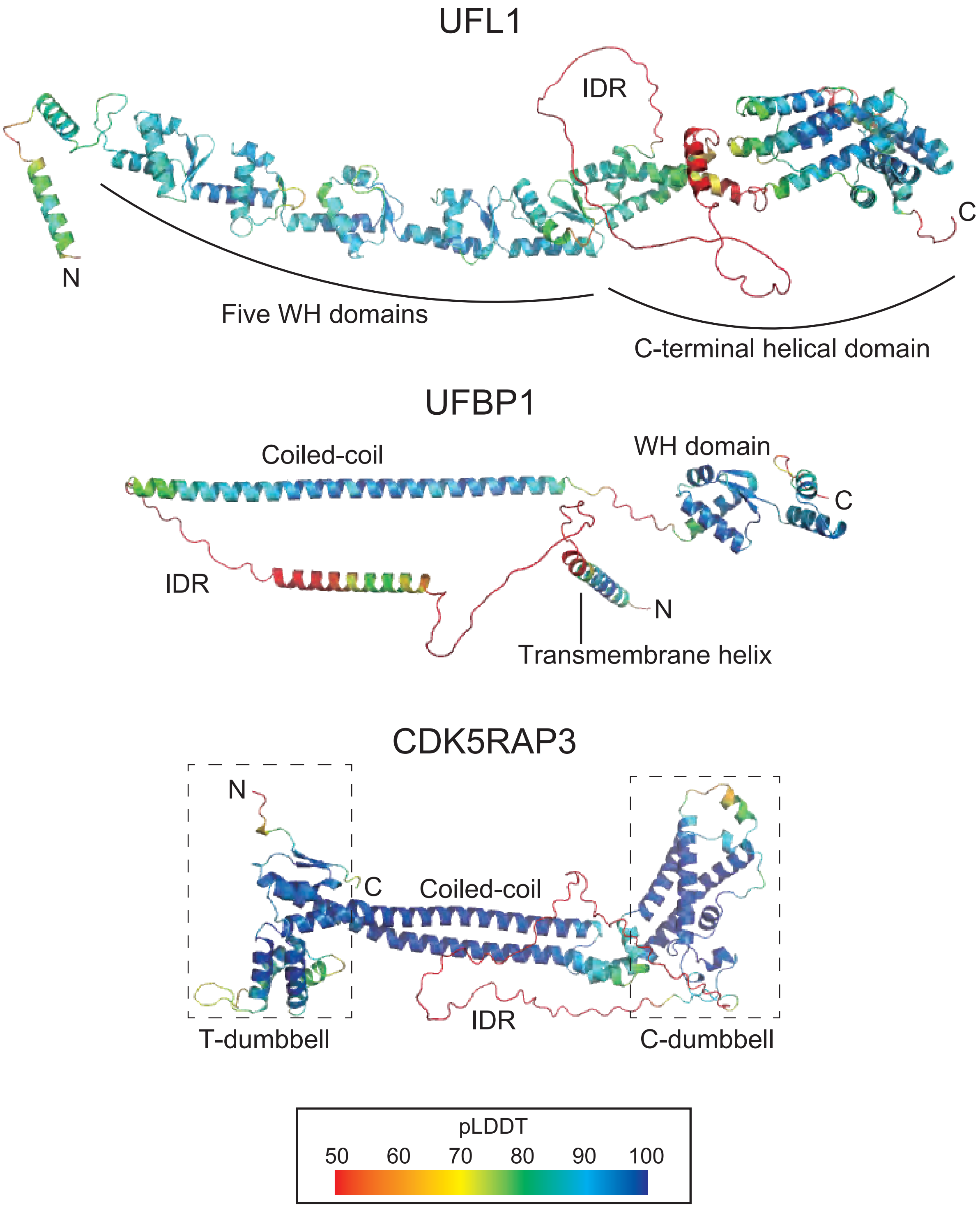


**Supplementary Figure 1**

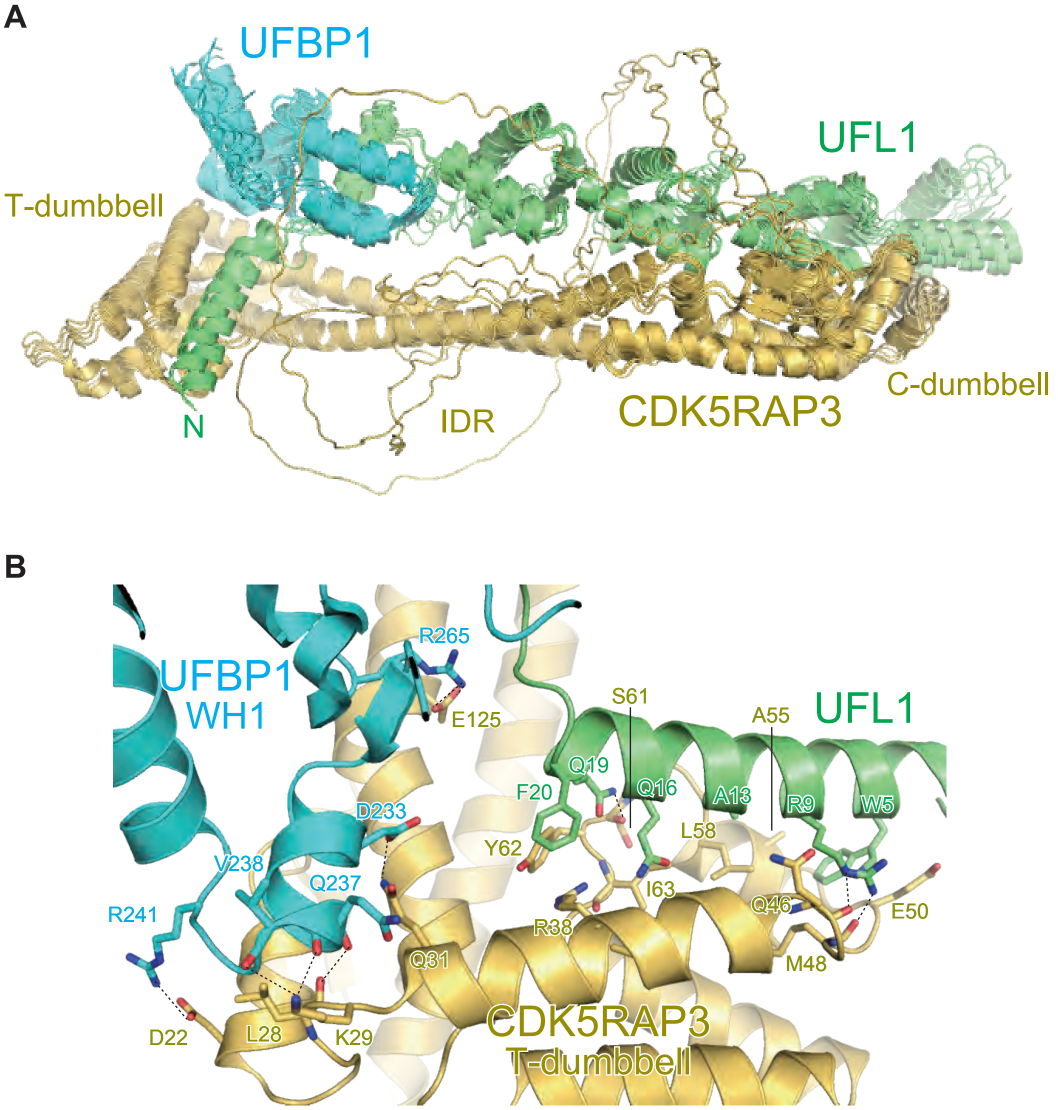


**Supplementary Figure 2**

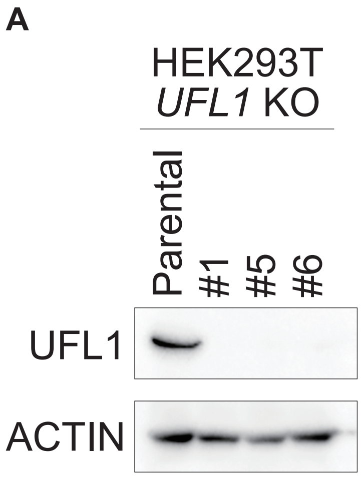


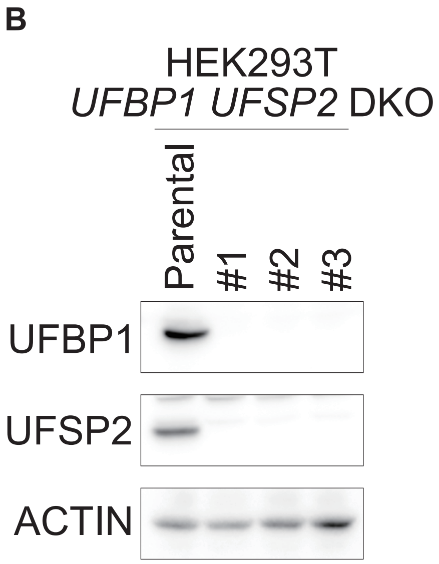


**Supplementary Figure 3**

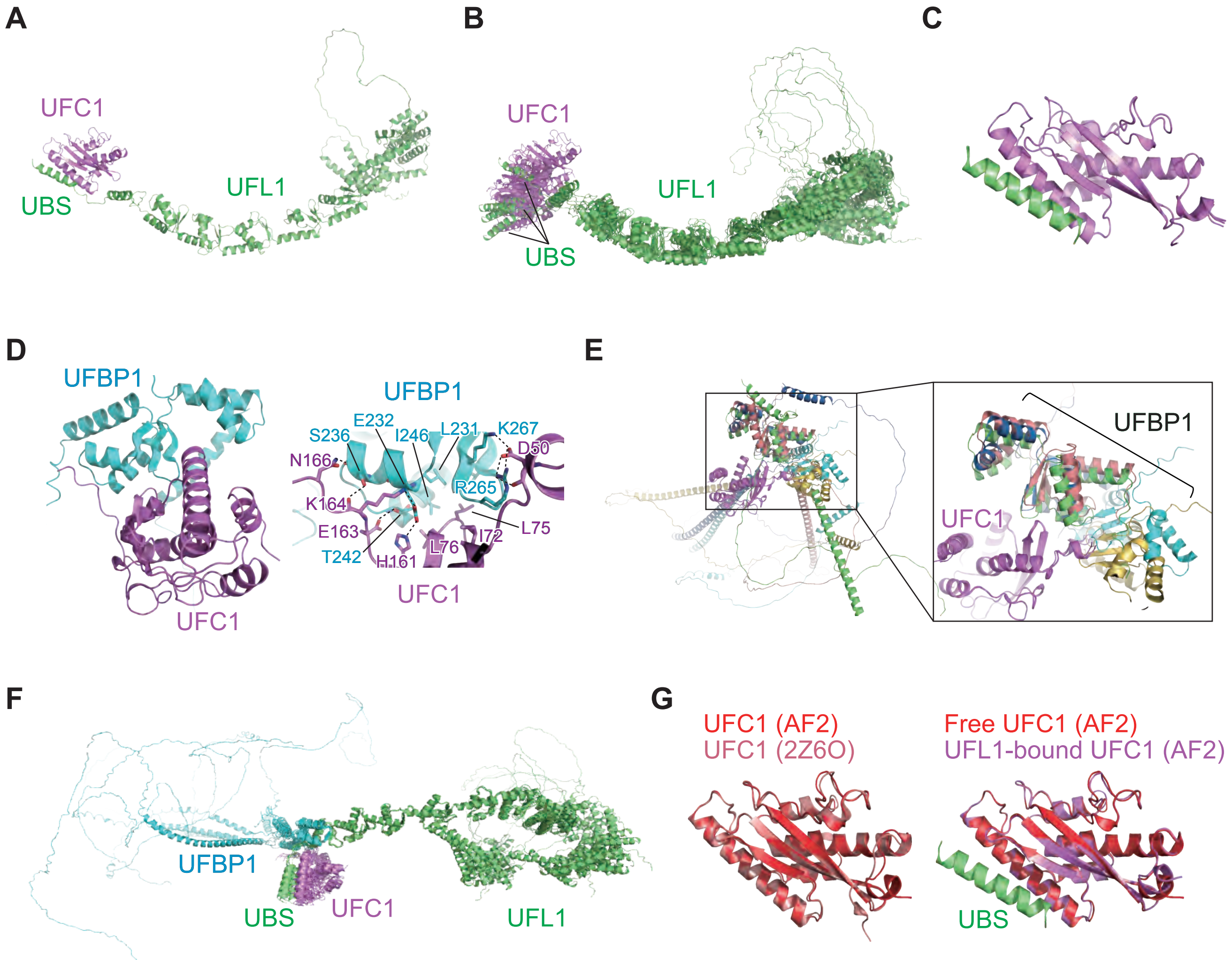


**Supplementary Figure 4**

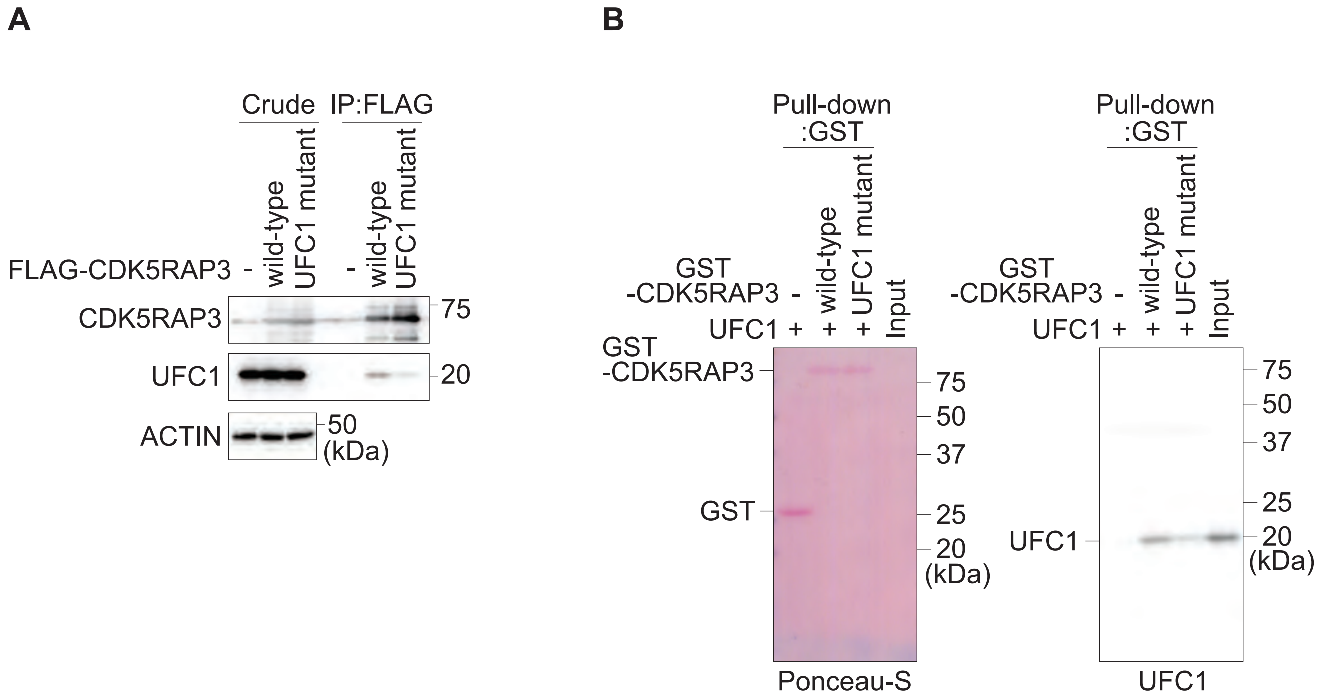


**Supplementary Figure 5**

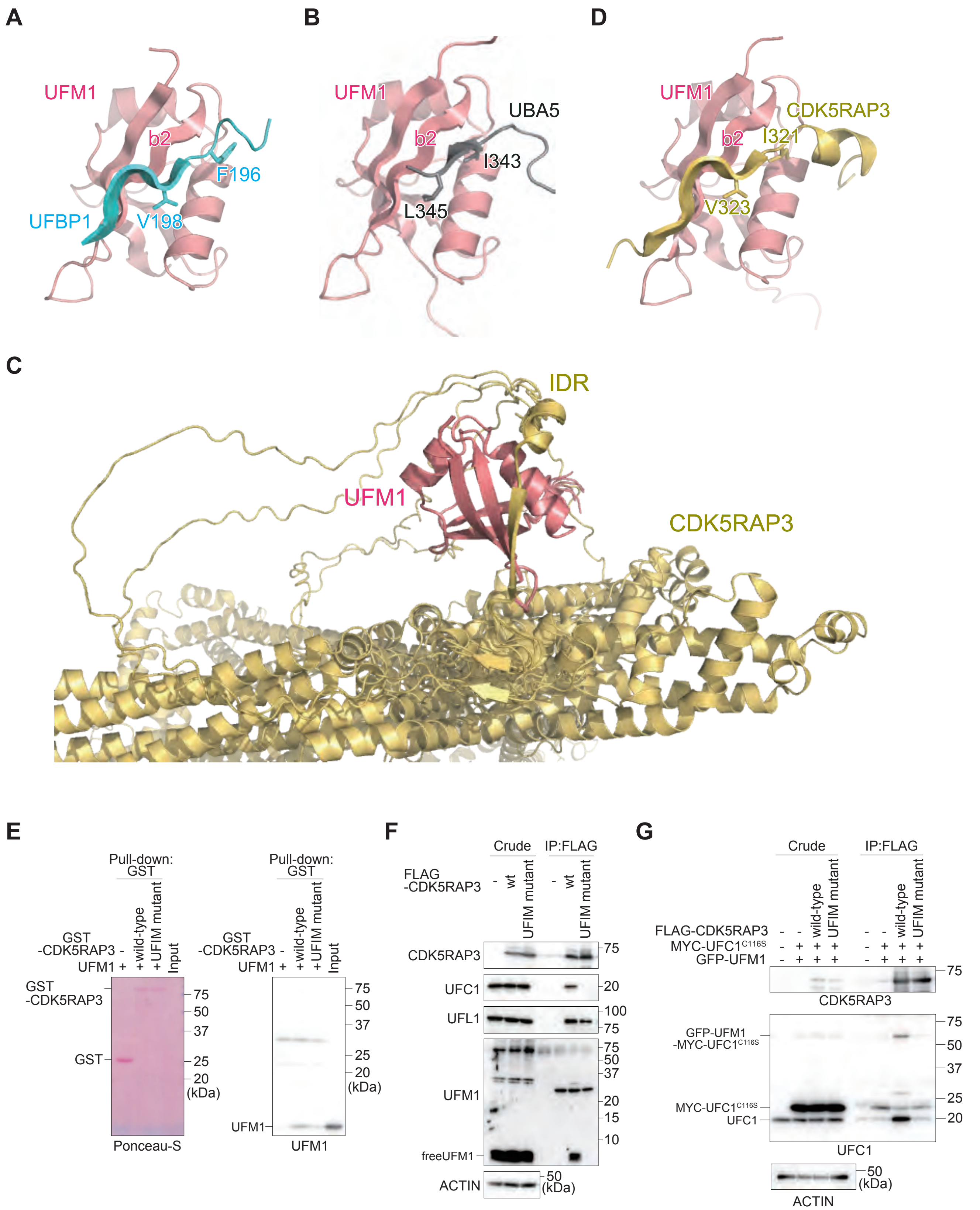


**Supplementary Figure 6**
